# Phage portal proteins counteract stringent-response–mediated restriction

**DOI:** 10.64898/2026.01.27.700999

**Authors:** Kristina Kronborg, Luokai Wang, Muriel Leandra Schicketanz, Kenn Gerdes, Yong Everett Zhang

## Abstract

Bacteria restrict viral replication not only through dedicated defense systems but also by entering global physiological states that limit cellular resources. How phages counter such host-imposed physiological barriers remains poorly understood. The stringent response (SR), mediated by the alarmones ppGpp and pppGpp, induces a growth-restrictive state that can act as a barrier to bacteriophage infection. Here we show that elevated alarmone levels delay T7-mediated host lysis, whereas (p)ppGpp-deficient cells are hypersensitive to infection, establishing alarmone signaling as a physiological constraint on phage replication. A systematic functional screen identifies the essential capsid portal protein Gp8 as a viral factor genetically linked to SR-dependent host physiology. Gp8 directly binds the alarmone synthetases RelA and SpoT, selectively inhibiting their synthetase activities *in vitro* and suppressing alarmone accumulation *in vivo*. Phages carrying interaction-defective portal mutations exhibit impaired replication, delayed lysis, and sustained (p)ppGpp elevation during infection, defects that are alleviated in SR-deficient hosts. Structural and mutational analyses reveal electrostatically mediated interfaces required for targeting RelA and SpoT. Portal proteins from diverse coliphages share similar structural features, interact with stringent-response enzymes, and display SR-linked phenotypes, indicating a broadly conserved viral strategy. Together, these findings identify phage portal proteins as a previously unrecognized class of viral counter-defense factors that target a central bacterial stress signaling pathway, revealing that essential structural virion components can moonlight as regulators of host stress physiology.

## Introduction

Bacteria employ multilayered defense networks to restrict viral infection, combining dedicated systems such as restriction-modification, CRISPR-Cas, abortive infection modules, and nucleotide-based immune signaling pathways with broader physiological states that limit viral replication^1-4^. Whereas many antiviral mechanisms rely on molecular recognition of invading phages, global physiological responses can impose resource constraints that indirectly restrict infection. Physiological growth arrest, a common outcome of abortive infection systems, limits phage spread by sacrificing infected cells^5^. How phages contend with such host-imposed physiological barriers remains incompletely understood.

One of the most ancient and conserved physiological responses is the stringent response (SR), mediated by the alarmones ppGpp and pppGpp^6,7^. Activation of the SR reprograms transcription and metabolism, suppressing ribosome biogenesis and nucleotide synthesis while promoting stress adaptation^8^. Although classically viewed as a stress-adaptation mechanism, the growth-restrictive state imposed by the SR creates an intracellular environment that is unfavorable for bacteriophage replication, suggesting that it can function as a physiological barrier to infection.

Consistent with this view, (p)ppGpp signaling influences bacteriophage infection outcomes. Cells lacking alarmone synthesis are hypersensitive to phage killing, whereas stationary-phase cells, characterized by elevated alarmone levels, often exhibit increased resistance^9-11^. Toxic Rel-family proteins have also emerged in multiple abortive infection systems^12-15^, underscoring the recurrent involvement of RSH-associated regulatory pathways in phage–host interactions. Conversely, certain phage-encoded factors, such as the alarmone-degrading enzyme ATD1^16^, directly modulate (p)ppGpp metabolism during infection. Together, these observations indicate that alarmone-linked stress networks are repeatedly engaged during defense and counter-defense processes.

Phages, in turn, have evolved diverse strategies to neutralize host defenses, including inhibitors of CRISPR-Cas systems, restriction enzymes, abortive infection pathways, and cyclic-nucleotide immune signaling modules^17,18^. Most characterized anti-defense factors target discrete molecular effectors. Whether phages also encode dedicated mechanisms to counteract global physiological restriction states, such as the SR, remains largely unexplored. Because the SR is governed by a small number of conserved enzymes, it represents a potentially vulnerable node for viral counter-defense.

In *Escherichia coli*, the SR is mediated primarily by the (p)ppGpp synthetases RelA and SpoT^19-23^. Elevated alarmone levels remodel transcription through direct interactions with RNA polymerase (RNAP)^24,25^, limiting biosynthetic capacity and slowing growth^26-28^. Although this reprogramming enhances stress survival, it may also constrain the metabolic demands of phage replication. Consistent with this, bacteriophage T7 encodes multiple inhibitors of host RNAP^11,29-31^, highlighting the importance of transcriptional control during infection. These observations raise the possibility that phages actively modulate SR signaling to optimize infection.

Here, we investigate the interplay between the SR and infection by bacteriophage T7. Using complementary genetic, biochemical, and phage mutational approaches, we identify the essential T7 capsid portal protein Gp8 as a direct inhibitor of (p)ppGpp synthesis. Gp8 physically interacts with RelA and SpoT, selectively suppressing their synthetase activities and reducing intracellular alarmone accumulation during infection. Disruption of this interaction impairs phage replication and delays host lysis. Portal proteins from diverse *E. coli* phages exhibit similar properties, revealing a broadly conserved strategy by which structural virion components moonlight as modulators of host stress physiology to promote infection.

## Results

### The stringent response limits bacteriophage infection

Bacterial stress responses reshape cellular physiology in ways that can influence susceptibility to viral infection. To test whether the stringent response (SR) affects bacteriophage replication, we examined infection by bacteriophage T7 in a (p)ppGpp⁰ *Escherichia coli* strain lacking both *relA* and *spoT*. Compared with wild-type cells, the alarmone-deficient host exhibited accelerated killing kinetics and more rapid lysis, as reflected by larger plaque sizes and increased plaque expansion rates (**Fig. 1a,b**). These results indicate that loss of alarmone signaling enhances phage infection, consistent with previous report^11^.

**Figure 1.**
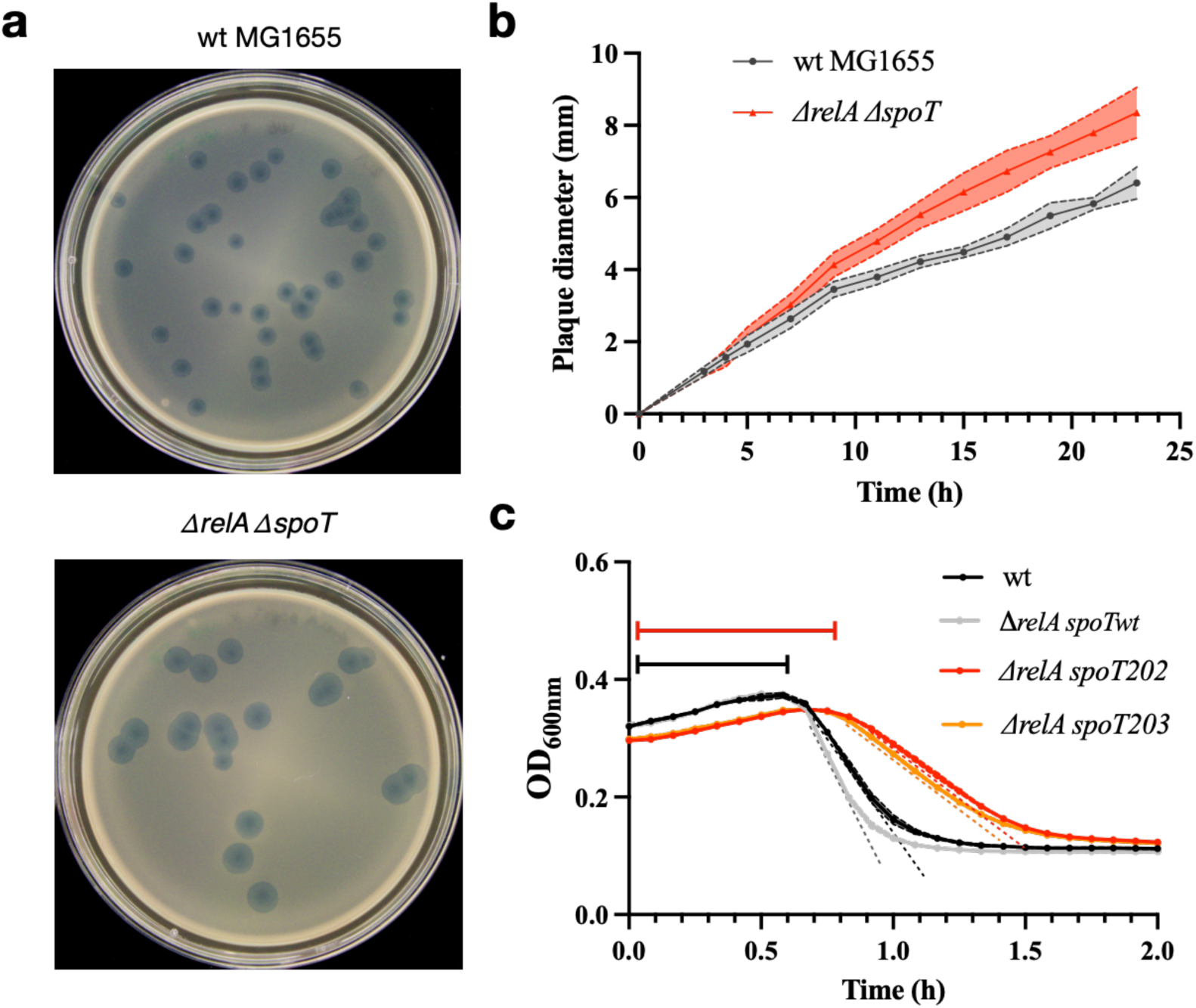
The stringent response restricts bacteriophage infection. **a**, Plaques of T7 phage after 24-hour infection of wild-type and (p)ppGpp⁰ (*ΔrelA ΔspoT*) *Escherichia coli* on LB medium. **b**, Quantification of plaque expansion kinetics for T7 infection of wild-type and (p)ppGpp⁰ (*ΔrelA ΔspoT*) cells, as in **a**. Data represent mean ± s.d. from five independent plaques per strain. **c**, Lysis kinetics of wild-type MG1655, *ΔrelA spoT*wt and high-(p)ppGpp strains (*ΔrelA spoT202* and *ΔrelA spoT203*) during T7 infection. Data represent mean values from independent biological replicates; shaded areas indicate s.d.. The lag phase preceding lysis and the lysis rate (slope) are indicated.

We next tested whether elevated alarmone levels have the opposite effect. Strains carrying *spoT* alleles that increase basal (p)ppGpp concentrations (*spoT202* and *spoT203*)^32,33^ displayed delayed lysis kinetics relative to *spoT* wild-type allele cells, characterized by prolonged lag phases prior to host population collapse and reduced lysis slopes (**Fig. 1c**). Across genetic backgrounds, higher intracellular alarmone levels were associated with slower infection progression.

Together, these results demonstrate that alarmone signaling establishes a growth-restrictive intracellular environment that limits bacteriophage replication. The SR therefore functions as a physiological barrier to infection, suggesting that successful phages must encode mechanisms to overcome this defense-associated state.

### T7 encodes factors that interfere with stringent-response-dependent host physiology

Because elevated alarmone levels restrict phage infection (**Fig. 1**), we reasoned that T7 may encode factors that counteract the SR**-**associated physiological state. Anti-defense systems often manifest as proteins that perturb host pathways required specifically under stress or nutrient-limited conditions. We therefore performed a systematic functional screen to identify T7 proteins that interfere with host growth under conditions in which (p)ppGpp signaling is required.

We cloned 53 T7 genes under the arabinose inducible control of the pBAD33 vector and assessed their effects on *E. coli* growth in rich (LB) and minimal media (M9) (**Fig. 2a**). Strikingly, 34 of 53 T7 proteins caused severe growth inhibition when expressed in minimal medium, whereas most exhibited little or no toxicity in rich medium (**Fig. 2b**; Supplementary **Fig. S1**). This conditional toxicity pattern suggests that T7 encodes numerous factors targeting host processes that become critical under nutrient limitation, a physiological context strongly influenced by alarmone signaling.

**Figure 2.**
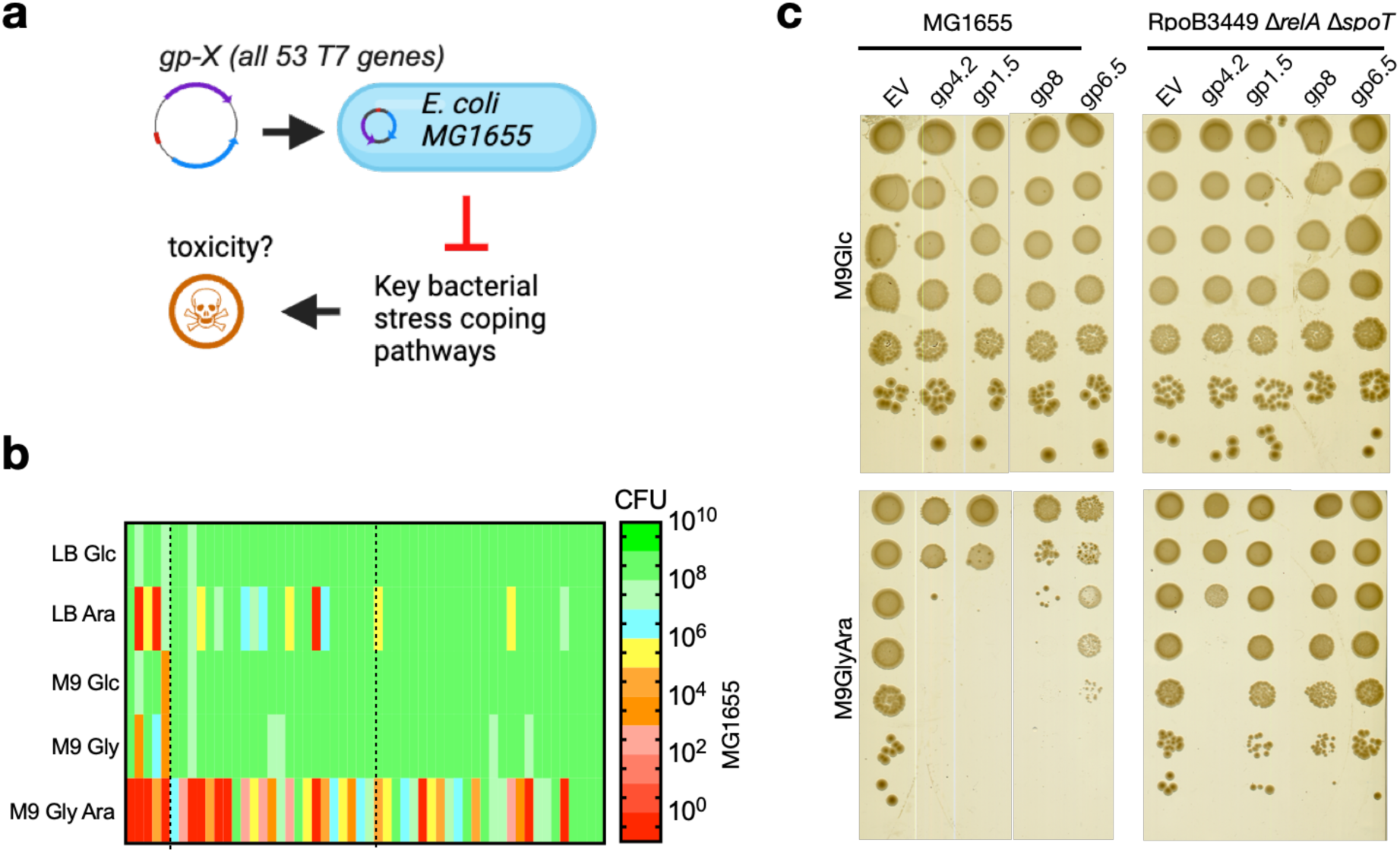
Identification of T7 proteins linked to stringent-response-dependent physiology. **a**, Schematic of the systematic testing of all 53 phage T7 genes into the inducible plasmid vector pBAD33 and their expression in *Escherichia coli* MG1655 to screen for phage proteins that impair bacterial growth under conditions requiring stringent-response function. (Created with BioRender.com) **b**, Growth of *E. coli* expressing individual T7 genes in rich and minimal media, quantified as CFU ml⁻¹ normalized to OD_600nm_. LB Glc and LB Ara are LB media supplemented with 0.2% glucose and 0.2% arabinose, respectively. M9 Glc, M9 Gly, and M9 Gly Ara denote M9 minimal media supplemented with 0.4% glucose, 0.4% glycerol, or 0.4% glycerol plus 0.2% arabinose, respectively. Original plate-based growth assays are shown in **Supplementary Fig. S1**. **c**, Suppression of T7 protein**-**induced growth defects in (p)ppGpp⁰ (*ΔrelA ΔspoT*) strains carrying stringent-like RNA polymerase allele (*rpoB*3449; RNAP^SR^). Serial dilutions of the indicated strains expressing gp1.5, gp4.2, gp6.5, gp8, or empty vector are shown.

To identify T7 proteins specifically linked to SR-dependent physiology, we next exploited RNA polymerase (RNAP) mutants with a stringent-like (RNAP^SR^) phenotype that bypass the requirement for (p)ppGpp during growth in minimal medium^34,35^. Expression of four T7 proteins, Gp1.5, Gp4.2, Gp6.5, and Gp8, showed reproducible suppression of toxicity in two independent RNAP^SR^ backgrounds (Supplementary **Fig. S2**-**S3**). Notably, suppression persisted in a (p)ppGpp⁰ strain carrying the RNAP^SR^ allele^34^ (**Fig. 2c)**, indicating that rescue does not require alarmone synthesis itself but rather depends on SR-regulated transcriptional programs.

Together, these results identify a subset of T7 proteins genetically linked to host SR pathways, consistent with the presence of viral factors that interfere with this defense-associated physiological state.

### The portal protein Gp8 directly targets the alarmone synthetases RelA and SpoT

The genetic link between a subset of T7 proteins and SR**-**dependent host physiology prompted us to test whether these candidates directly target components of the alarmone signaling machinery. We therefore examined physical interactions between Gp1.5, Gp4.2, Gp6.5, and Gp8 and the central SR regulators RelA, SpoT, and RNAP (i.e., the ω subunit RpoZ) using a bacterial two-hybrid assay. Among the proteins tested, the T7 portal protein Gp8 showed robust and reproducible interactions with both RelA and SpoT (**Fig. 3a**), whereas the other candidates did not display consistent interactions.

**Figure 3.**
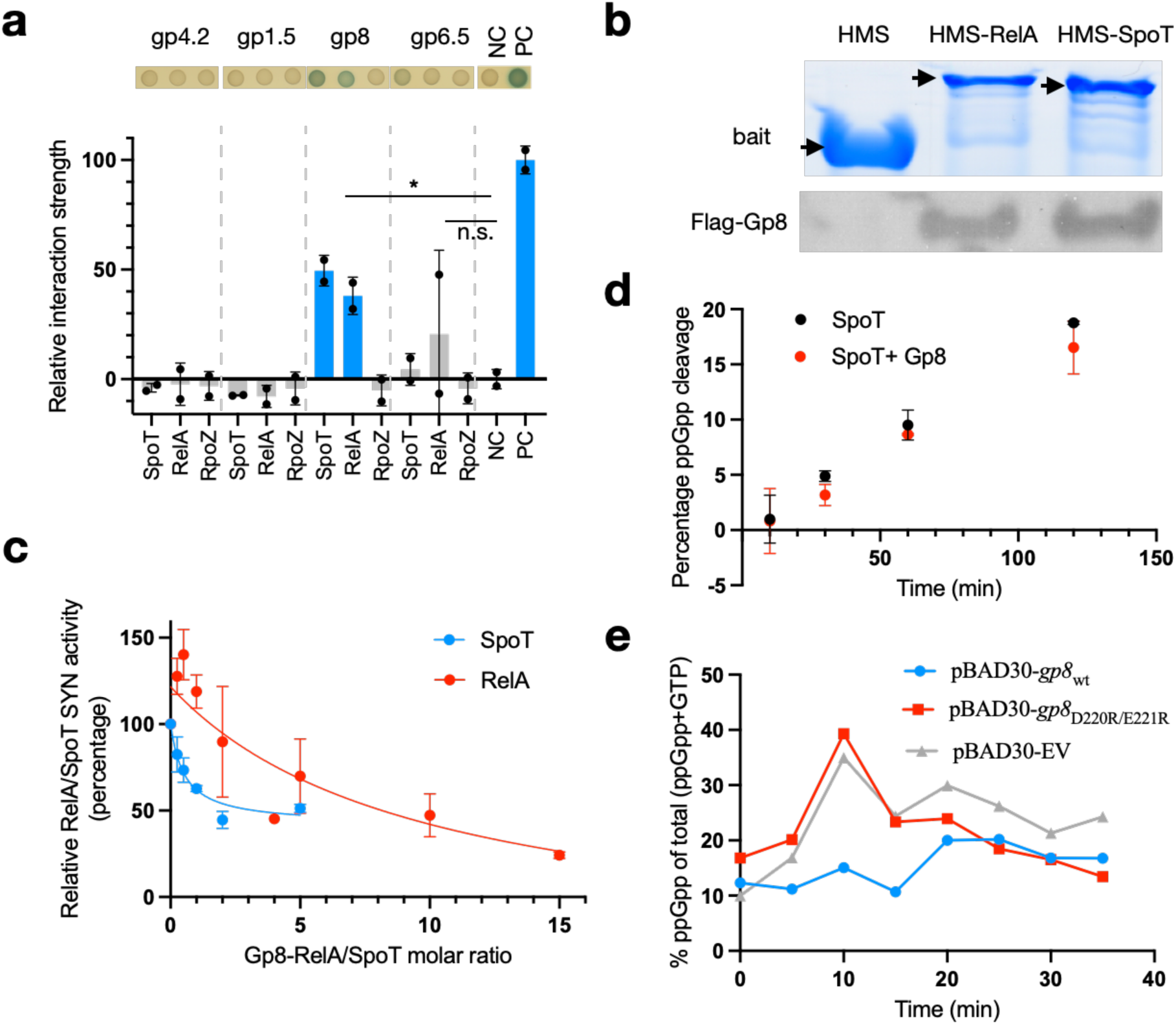
The portal protein Gp8 inhibits the alarmone synthetases RelA and SpoT. **a**, Bacterial two-hybrid assays testing interactions between T7 proteins and RelA, SpoT, or RpoZ (the ω subunit of RNA polymerase). Data represent mean ± s.d. from two independent experiments. NC and PC denote negative and positive controls, respectively. * indicates *P* < 0.05 by Student’s *t*-test. Representative spots are shown above the corresponding bars. **b**, Pull-down assays showing co-purification of Flag**-**Gp8 with His**-**MBP**-**SUMO (HMS)**-**tagged RelA or SpoT, or with the HMS tag alone. Arrows indicate bands corresponding to the bait proteins. **c**, *In vitro* (p)ppGpp synthetase activity of RelA and SpoT in the presence of increasing molar ratios of Gp8. Data represent mean ± s.d. from three independent experiments. **d**, SpoT hydrolase activity measured in the absence or presence of Gp8. Data represent mean ± s.d. from two independent experiments. **e**, Intracellular ppGpp levels, normalized to total (ppGpp + GTP), following Gp8 expression from the pBAD30 vector, measured by thin-layer chromatography. EV, empty vector control.

We validated these associations biochemically by co-purification assays using Flag-tagged Gp8 and His**-**MBP**-**SUMO(HMS)-tagged RelA or SpoT. In both cases, Gp8 specifically copurified with the alarmone synthetases, confirming direct physical interaction *in vitro* (**Fig. 3b**). These results identify Gp8, an essential structural component of the T7 capsid portal vertex, as a viral protein that directly engages the core enzymatic machinery responsible for (p)ppGpp synthesis.

We next asked whether Gp8 binding affects the catalytic activities of RelA and SpoT. *In vitro* enzymatic assays revealed that Gp8 potently inhibited the synthetase activities of both enzymes in a dose-dependent manner (**Fig. 3c**). In contrast, Gp8 had no detectable effect on the hydrolase activity of SpoT (**Fig. 3d**), indicating that the portal protein selectively suppresses alarmone synthesis rather than globally perturbing SpoT function. Consistent with this, expression of Gp8 *in vivo* suppressed intracellular (p)ppGpp accumulation relative to control conditions (**Fig. 3e** and Supplementary **Fig. S3**).

Together, these findings demonstrate that the phage portal protein Gp8 directly binds and inhibits the alarmone synthetases RelA and SpoT, revealing a mechanism by which T7 suppresses the SR pathway.

### Electrostatic interfaces on the portal ring mediate targeting of RelA and SpoT

Gp8 assembles into a dodecameric ring that forms the portal vertex of the T7 capsid, with exterior surfaces enriched in charged residues (**Fig. 4a**). To define the regions of Gp8 required for interaction with RelA and SpoT, we generated domain truncations and assessed binding using the bacterial two-hybrid assay. Deletion of either the N- or C-terminal regions of Gp8 abolished interactions with both RelA and SpoT, whereas removal of the clip domain reproducibly enhanced interaction signals. Further deletion extending into the stem domain eliminated binding entirely (**Fig. 4b**). These results indicate that an intact portal ring architecture is required for targeting the alarmone synthetases and suggest that the clip motif does not itself contribute to binding but instead modulates access to the interaction surface.

**Figure 4.**
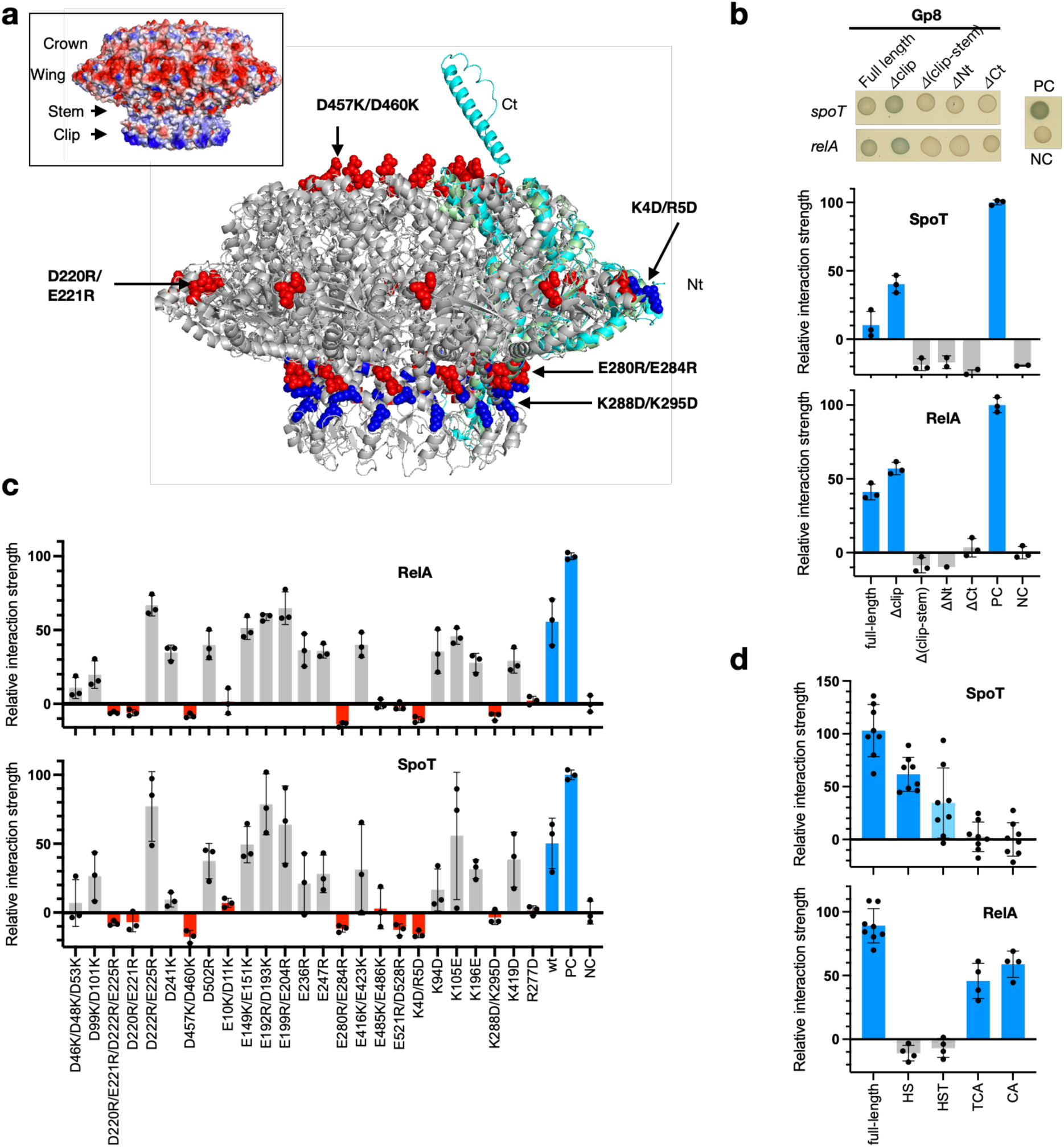
Portal surface charge interfaces mediate interaction with RelA and SpoT. **a**, Structure of the T7 Gp8 portal ring (grey ribbon model; PDB 7EY6). An AlphaFold model of the full-length Gp8 monomer (cyan) is overlaid with a single monomer from the portal ring structure (green) to indicate the positions of the N- and C-terminal regions, including the mutated K4 and R5 residues. Residues whose mutation abolished interaction with RelA and SpoT are shown as spheres and labeled. Negatively charged residues are shown in red and positively charged residues in blue. Inset: electrostatic surface representation of the Gp8 portal ring. **b**, Bacterial two-hybrid assays assessing interactions between Gp8 truncation variants and full-length RelA or SpoT. Δclip, deletion of residues 299**-**330; Δ(clip**-**stem), deletion of residues 269**-**355; ΔNt, deletion of residues 6**-**42; ΔCt, deletion of residues 484**-**543. NC and PC denote negative and positive controls, respectively. Bar plots show quantification of relative interaction strengths, normalized to the positive control (PC). NC corresponds to cells carrying empty vectors. **c**, Quantification of relative interaction strengths between Gp8 point mutants and RelA or SpoT in bacterial two-hybrid assays, normalized to the positive control (PC). NC indicates negative control cells containing empty vectors. WT denotes interactions with the respective wild-type RelA or SpoT proteins. Data represent means from three independent replicates. Red bars indicate Gp8 mutants that completely lost interaction with both RelA and SpoT. Representative raw spot images are shown in Supplementary **Fig.** S4. **d**, Quantification of relative interaction strengths between Gp8 and truncated variants of RelA or SpoT, normalized to the respective full-length proteins. Representative data from at least four independent replicates are shown. Domain abbreviations: HST, hydrolase**-**synthetase**-**TGS domains; HS, hydrolase**-**synthetase domains; TCA, TGS**-**CC**-**ACT domains; CA, CC**-**ACT domains. Precise domain boundaries of the RelA and SpoT truncations, along with representative raw spot images, are provided in Supplementary **Fig.** S4.

Notably, the clip motif is enriched in positively charged residues, whereas the remainder of the portal exterior is predominantly electronegative (**Fig. 4a**). The observation that deletion of a positively charged element enhances binding suggests that RelA and SpoT preferentially engage negatively charged surfaces on the assembled portal ring and that the clip motif may partially shield or sterically constrain these interfaces in the intact protein. We therefore tested whether electrostatic interactions underlie RelA/SpoT binding by generating 43 charge-reversal mutants targeting clusters of surface-exposed acidic and basic residues.

Approximately half of the charge-reversal mutants exhibited substantial loss of interaction with RelA, and the same substitutions similarly impaired SpoT binding (**Fig. 4c** and Supplementary **Fig. S4a-e**). Several mutations - including D220R/E221R, D457K/D460K, E280R/E284R, K4D/R5D, and K288D/K295D - abolished interaction entirely, identifying discrete charged surface patches (**Fig. 4a)** that are jointly required for targeting both enzymes. These results support a model in which RelA and SpoT recognize a defined electrostatic landscape on the portal ring.

To define the corresponding interfaces on the host proteins, we performed domain mapping of RelA and SpoT. Removal of the N-terminal catalytic domains of SpoT eliminated its interaction with Gp8, whereas truncation of the C-terminal regulatory domains of RelA disrupted binding (**Fig. 4d** and Supplementary **Fig. S4f,g**). Thus, Gp8 engages RelA and SpoT through distinct domain interfaces, while relying on a common set of electrostatically defined surfaces on the portal ring.

Together, these data demonstrate that the portal protein uses a spatially organized pattern of surface charges to target the alarmone synthesis machinery, supporting the idea that suppression of the SR reflects a specific, evolved molecular interaction rather than a secondary consequence of portal assembly.

### Suppression of alarmone signaling by Gp8 is required for efficient phage infection

To determine whether Gp8-mediated targeting of RelA and SpoT contributes to phage fitness, we generated T7 phages carrying portal mutations that disrupt interaction with the alarmone synthetases. Using recombineering, we isolated mutant phages bearing substitutions D220R/E221R (interaction-defective) and E10K/D11K (partially defective), and confirmed impaired binding to RelA and SpoT by pull-down assays (**Fig. 5a**).

**Figure 5.**
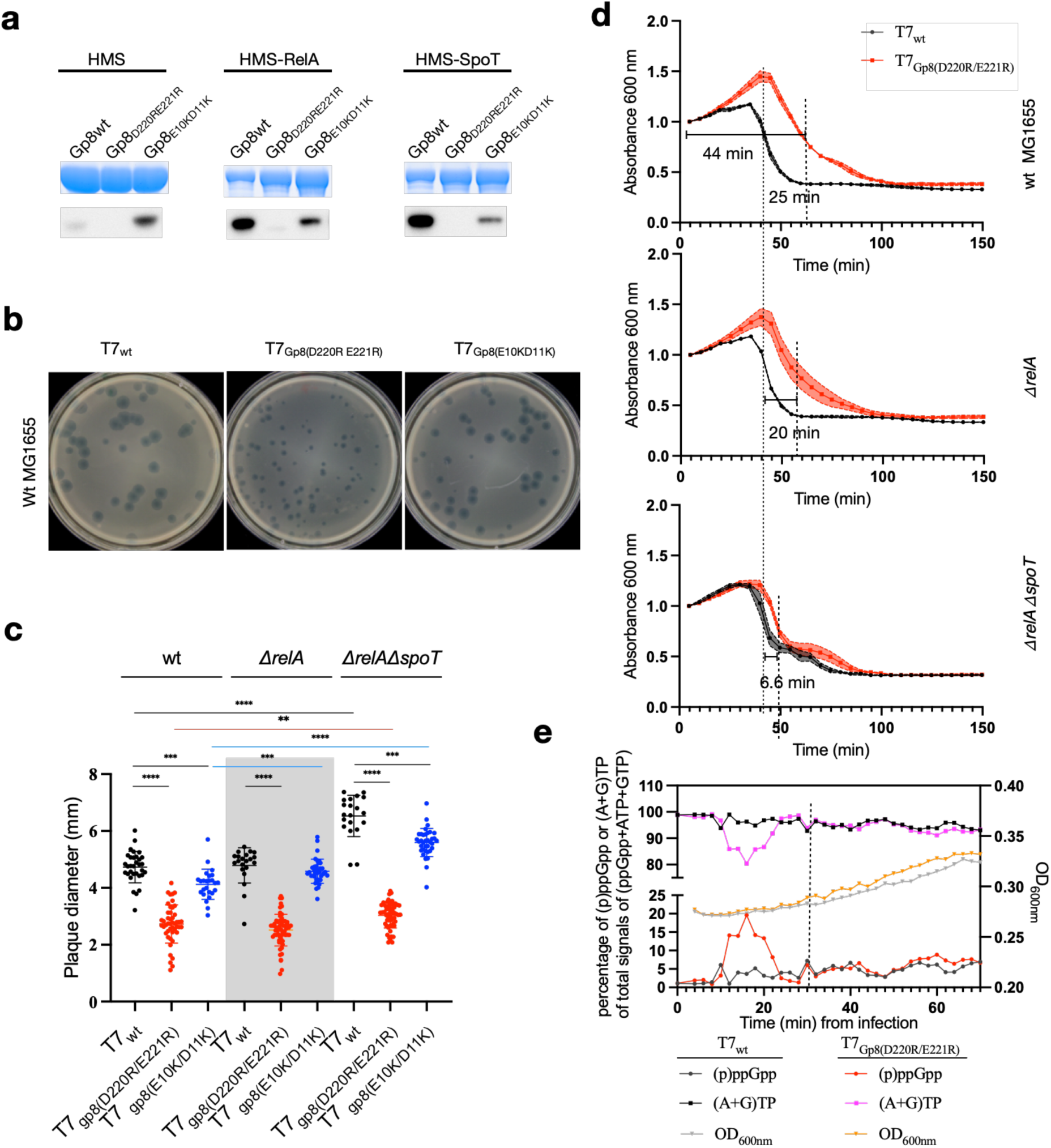
Portal-mediated suppression of alarmone signaling is required for efficient infection. **a**, Pull-down assays using HMS-tag, HMS-RelA, or HMS-SpoT as bait with Flag-tagged wild-type (WT) or mutant Gp8 proteins. Top: SDS-PAGE analysis of the bait proteins; bottom: immunoblot detection of co-purified Flag-Gp8 using an anti-Flag antibody. **b**, Representative plaque morphologies of WT or mutant T7 phages infecting WT *E. coli* MG1655 cells, imaged 18 h post infection. **c**, Quantification of plaque sizes corresponding to panel **b** for WT and mutant T7 phages infecting WT, Δ*relA*, or (p)ppGpp⁰ (Δ*relA* Δ*spoT*) strains. Data are shown as scatter plots. **, ***, and **** denote Student’s *t*-test *P* < 0.01, 0.001, and 0.0001, respectively. (**d**) Liquid infection assays of WT or mutant T7 phages infecting WT, Δ*relA*, or (p)ppGpp⁰ (Δ*relA* Δ*spoT*) strains. Dashed lines indicate the time points at which 50% of the total OD_600nm_ decrease associated with lysis had occurred; corresponding durations and differences are indicated. **e**, Quantification of ATP, GTP, and (p)ppGpp levels in radiolabeled *E. coli* MG1655 cells following infection with WT or mutant T7 phage. OD_600nm_ values of the host cultures are plotted on the secondary (right) *y*-axis. Experiments were performed in MOPS medium supplemented with 0.2 mM K₂HPO₄ and 0.2% (w/v) glucose. Representative thin-layer chromatography plates are shown in Supplementary **Fig. S7**. The first cycle of cell lysis (indicated by a dashed line) occurred approximately 30 min post infection.

In plaque assays on wild-type *E. coli*, the interaction-defective mutant formed significantly smaller plaques than T7_wt_, whereas the partially defective mutant displayed an intermediate phenotype (**Fig. 5b,c**). These defects were alleviated in hosts lacking SR signaling: plaque sizes increased in Δ*relA* cells for the partially defective mutant and in (p)ppGpp⁰ cells for all phages tested, indicating that impaired infection is linked to the host alarmone pathway.

We next quantified infection dynamics in liquid culture at a multiplicity of infection (MOI) of 0.1. Compared with T7_wt_, the interaction-defective mutant exhibited a marked delay in host cell lysis during infection of wild-type cells (**Fig. 5d**). This delay was substantially reduced in Δ*relA* and nearly abolished in (p)ppGpp⁰ strains, demonstrating that elevated alarmone levels are responsible for much of the infection defect (**Fig. 5d**). This is consistent with the prior observation that hosts with genetically elevated basal (p)ppGpp levels displayed delayed lysis (**Fig. 1b**), supporting an inverse relationship between alarmone abundance and infection efficiency.

To directly monitor alarmone dynamics during infection, we radiolabeled intracellular nucleotide pools and quantified (p)ppGpp levels over time. Because complex media such as LB interfere with nucleotide labeling, these experiments were performed in defined MOPS medium. Importantly, the relative infection phenotypes observed in LB were preserved under these conditions, with wild-type T7 lysing *E. coli* more rapidly than the interaction-defective mutant (Supplementary **Fig. S7a**).

Infections were performed at the same MOI of 0.1, such that population-wide lysis requires at least two rounds of infection, and alarmone dynamics are most clearly resolved during the first, relatively synchronized infection cycle. During this initial cycle, infection with the interaction-defective T7 mutant resulted in sustained elevation of (p)ppGpp prior to the onset of lysis, whereas alarmone levels were markedly reduced during wild-type T7 infection (**Fig. 5e** and Supplementary **Fig. S7b,c**). Elevated (p)ppGpp levels coincided with pronounced depletion of GTP and ATP pools, consistent with SR-associated repression of nucleotide biosynthesis^36-39^. At later time points corresponding to subsequent infection cycles and final population collapse, differences in alarmone levels between wild-type and mutant infections were diminished, likely reflecting increased heterogeneity in infection timing following the first lysis event.

Together, these results indicate that portal-mediated suppression of alarmone signaling promotes efficient T7 infection and that disruption of the Gp8-RelA/SpoT interaction leads to sustained SR activation and delayed infection progression.

### Portal-mediated suppression of the stringent response is conserved among coliphages

Gp8 is an essential structural component of the phage capsid and is widely conserved among tailed bacteriophages. To determine whether portal-mediated targeting of the SR represents a broader viral strategy, we examined portal proteins from phylogenetically diverse *E. coli* phages in the BASEL collection^40^. Structural predictions indicated that many of these portal proteins assemble into ring-like complexes with predominantly electronegative exterior surfaces, similar to that of T7 Gp8 (**Fig. 6a**; Supplementary Document 2).

**Figure 6.**
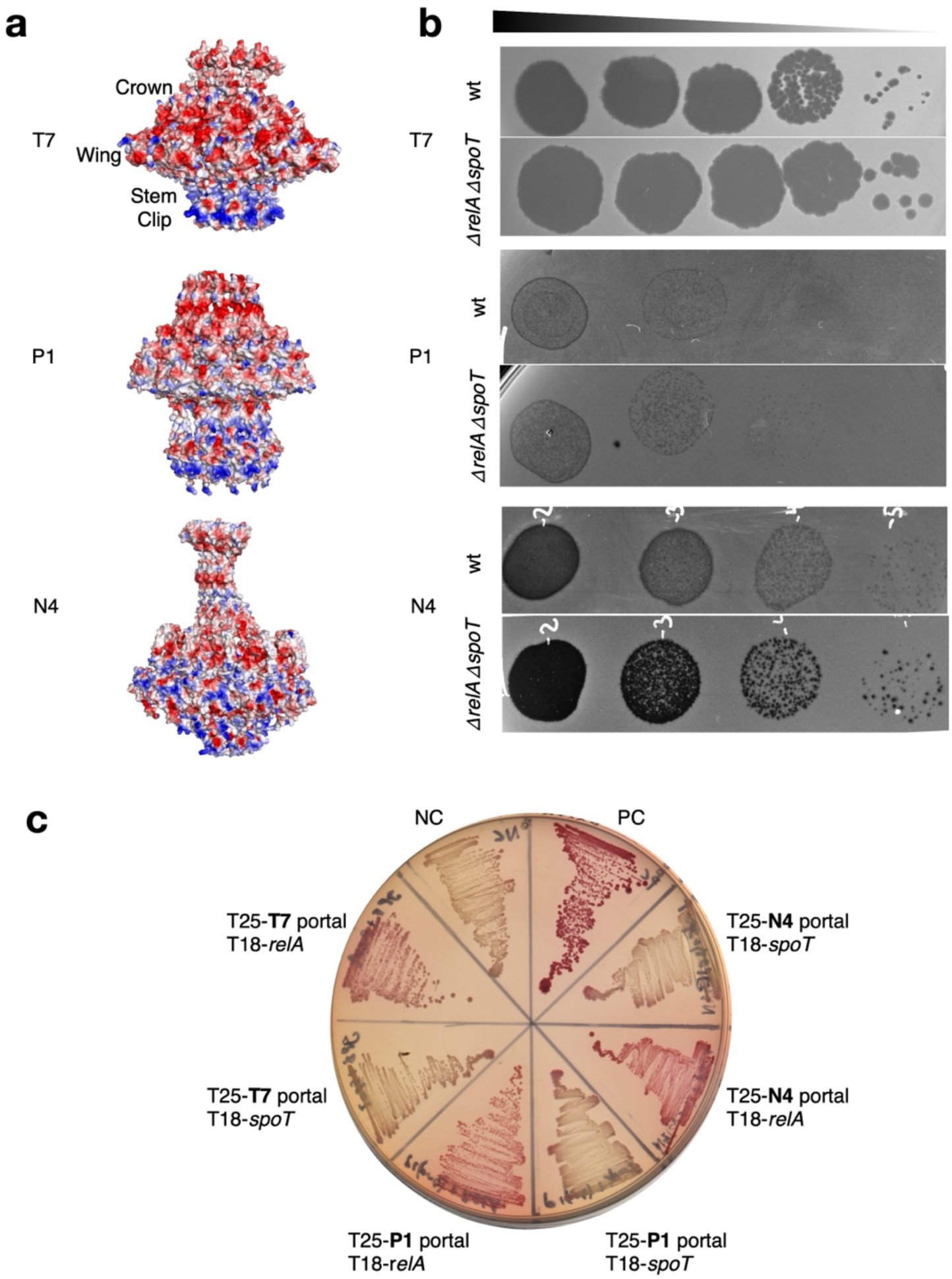
Portal-mediated suppression of the stringent response is conserved among coliphages. **a**, Predicted structural models of portal proteins from representative coliphages (T7, P1, N4), colored by electrostatic surface potential, highlighting conserved charge patterning across portals. Due to computational limitations, only half of the portal ring is displayed. **b**, Infection efficiency of representative phages assessed by tenfold serial dilution spotting on wild-type and (p)ppGpp⁰ (*ΔrelA ΔspoT*) *E. coli* hosts. **c**, Bacterial two-hybrid assays testing interactions between portal proteins from T7, P1, and N4 and the alarmone synthetases RelA and SpoT, visualized on MacConkey indicator plates.

Consistent with a general role for the stringent response in restricting infection, many of these phages infected (p)ppGpp⁰ cells more efficiently than wild-type hosts, producing larger or clearer plaques or increased plaque-forming units (**Fig. 6b**; Supplementary Document 2). Notably, portal proteins from these phages also displayed a conserved pattern of surface charge segregation, with negatively charged exterior regions juxtaposed to discrete positively charged elements (**Fig. 6a**), suggesting that this electrostatic organization may be a common feature associated with SR sensitivity.

To test whether portal proteins with this charge organization directly engage the alarmone synthesis machinery, we focused on representatives from two unrelated coliphages, P1 and N4. Although these phages differ substantially in genome organization, replication strategy, and virion architecture, their portal proteins form dodecameric ring assemblies analogous to that of T7 Gp8. In both cases, the portal proteins displayed a nonuniform surface charge distribution characterized by a positively charged clip-like region juxtaposed to a predominantly electronegative exterior surface (**Fig. 6a**). Expression of the P1 and N4 portal proteins was toxic in *E. coli* under conditions requiring SR function, and this toxicity was partially suppressed by RNAP^SR^ alleles (Supplementary **Fig. S8**). In bacterial two-hybrid assays, both portal proteins interacted with RelA and SpoT (**Fig. 6c**).

Together, these results demonstrate that portal proteins from evolutionarily distinct coliphages share conserved surface charge features and the capacity to interact with the alarmone synthesis machinery, indicating that portal-mediated modulation of stringent-response signaling is not unique to bacteriophage T7.

## Discussion

### A conserved role for phage portal proteins in repression of the stringent response

Together, these findings indicate that the SR can function not only as a stress adaptation pathway but also as a physiological barrier that constrains phage replication. While the SR has long been recognized for its role in bacterial survival under nutrient limitation and environmental stress, its capacity to globally suppress biosynthesis, transcription, and nucleotide availability also creates an intracellular environment that is unfavorable for viral replication. In contrast to dedicated antiviral defense systems that rely on molecular recognition of invading phages, the SR represents a distinct mode of host defense: rather than detecting phage components directly, it enforces a system-level physiological state that indirectly restricts viral growth by limiting access to essential cellular resources. Such global restriction mechanisms may be particularly effective against diverse phages, as they do not depend on specific recognition motifs that can be readily mutated or evaded.

### A biphasic model for SR modulation during T7 infection

Based on our data, we propose a biphasic model for the role of (p)ppGpp during T7 infection (**Fig. 7**). Early in infection, phage invasion and host takeover trigger stress responses that activate (p)ppGpp synthesis. At this stage, suppression of host metabolism, particularly inhibition of host RNAP, is advantageous to the phage, as it limits competing cellular processes and facilitates the transition to phage-directed transcription^41^. Notably, (p)ppGpp preferentially inhibits *E. coli* RNAP while having little effect on T7 RNAP^42,43^, allowing phage transcription to proceed efficiently. This effect is reinforced by multiple T7-encoded inhibitors of host RNAP (Gp2^29,31^, Gp5.7^11^, Gp0.7^30^), underscoring the importance of early host transcriptional shutdown for productive infection^41^.

**Figure 7.**
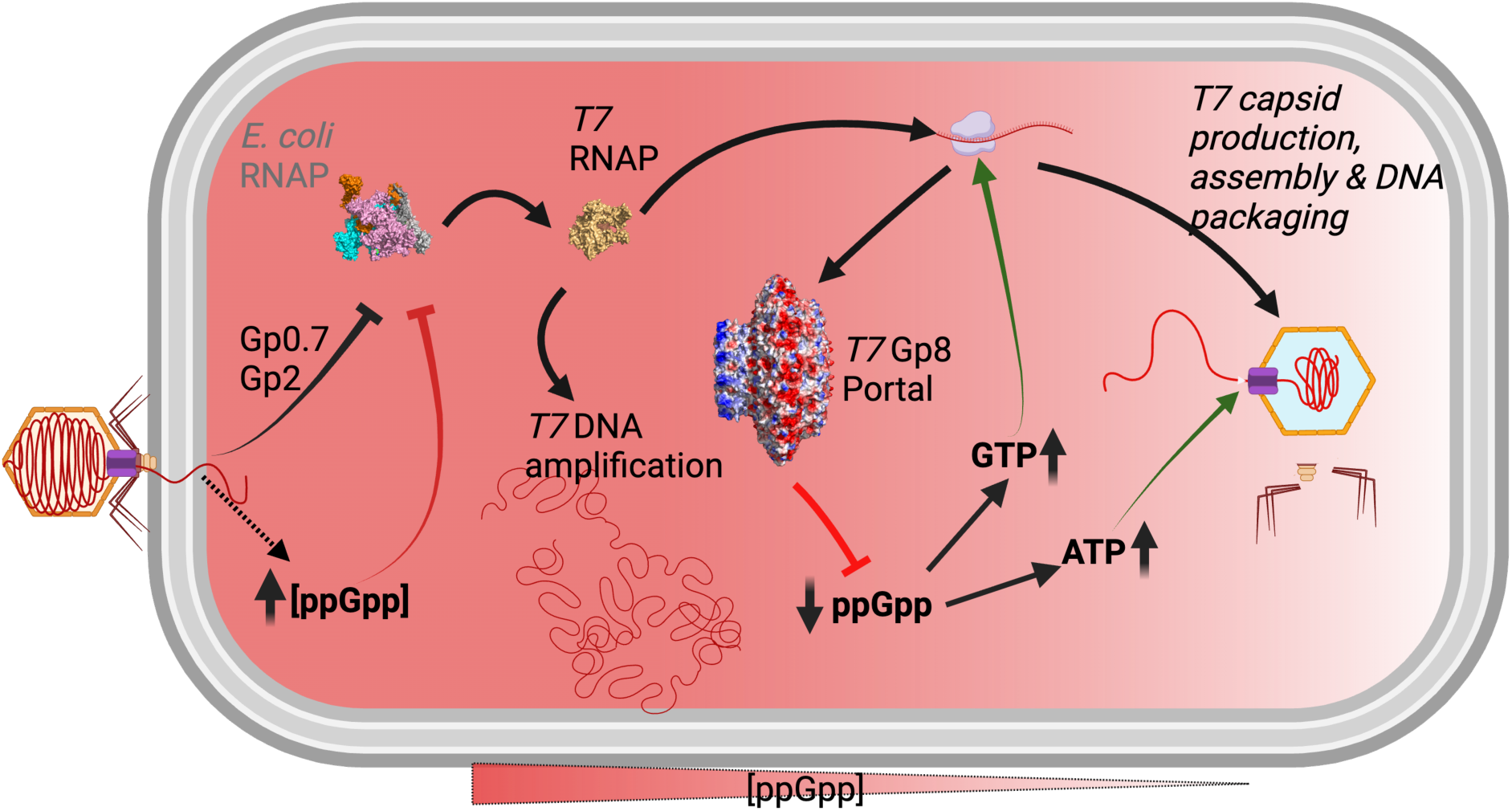
Biphasic model for portal-mediated modulation of the stringent response during phage infection. During early infection, phage entry and host takeover are associated with activation of the stringent response and accumulation of (p)ppGpp, which suppresses host transcription and contributes to the transition toward phage-directed gene expression. As infection progresses, the structural portal protein Gp8 accumulates and engages the alarmone synthetases RelA and SpoT, selectively inhibiting (p)ppGpp synthesis. Suppression of alarmone signaling restores nucleotide availability and biosynthetic capacity required for late-stage viral protein synthesis, genome packaging, and lysis. Portal-mediated targeting of stringent-response enzymes is conserved among coliphages, indicating that modulation of alarmone signaling represents a broadly conserved viral counter-defense strategy. (Created with BioRender.com)

As infection progresses, however, the physiological requirements of the phage shift. Late stages of infection involve massive synthesis of structural proteins and energy-intensive genome packaging by the terminase**-**portal complex, processes that place high demands on cellular amino acids and nucleotide pools^44,45^. Uncontrolled (p)ppGpp accumulation becomes detrimental at this stage, as it depletes ATP and GTP and further suppresses their biosynthesis^36-39^. Our data indicate that Gp8-mediated inhibition of RelA and SpoT alleviates this constraint by suppressing (p)ppGpp levels prior to lysis, thereby restoring nucleotide availability and enabling efficient virion assembly and release.

In this framework, the portal protein functions as a molecular gatekeeper that coordinates the transition from early host metabolic repression to late-stage resource mobilization, ensuring optimal timing of lysis and maximal phage yield. This biphasic control highlights how phages not only tolerate host defense-associated stress responses but actively repurpose them to optimize different phases of their replication cycle.

### Evolutionary implications of portal proteins as anti-defense factors

The identification of a regulatory role for the phage portal protein is evolutionarily striking. Portal proteins are among the most conserved and structurally constrained components of tailed phages, forming the central channel through which viral DNA is packaged and later delivered into the host. Their essential architectural role would be expected to limit sequence plasticity and functional diversification. Nonetheless, our data reveal that portal proteins have evolved spatially organized, electrostatically defined surface features capable of engaging host regulatory enzymes and modulating global signaling pathways. This suggests that strong selective pressure has favored coupling of genome delivery and packaging machinery with mechanisms that suppress host stress physiology. Such multifunctionality may ensure that evasion of host physiological restriction is tightly coordinated with late-stage replication demands, minimizing the risk that host defenses interfere with energetically costly assembly and packaging processes. More broadly, these findings expand the conceptual scope of phage anti-defense strategies by demonstrating that essential structural virion components - not only small, dedicated inhibitors - can moonlight as modulators of host stress physiology.

### Multifunctional phage proteins and conditionally essential host processes

The identification of a regulatory role for the portal protein highlights a broader principle in phage biology: the evolution of multifunctional proteins that target host processes essential under specific physiological conditions. Such functional layering is not unique to Gp8. For example, T7 lysozyme (Gp3.5) not only mediates host cell lysis but also regulates T7 RNAP to coordinate transcription and replication^46,47^. These dual-purpose proteins enable phages to maximize functional output from compact genomes.

Consistent with this idea, our functional screens revealed numerous T7 proteins that are toxic specifically under nutrient-limited conditions but not in rich media. This conditional toxicity suggests that phages encode factors targeting host pathways that become essential during stress, a context frequently encountered in natural and clinical environments but rarely captured in standard laboratory assays. As bacteria in nature often reside in nutrient-poor or otherwise stressful niches, phage strategies that exploit stress-adapted host physiology are likely widespread and ecologically relevant.

### Broader implications and future directions

The identification of RSH-family toxins such as CapRel in abortive infection systems^12-15^ further underscores the recurrent involvement of alarmone-linked regulatory pathways in phage–host interactions. In these systems, perturbation of translational and nucleotide homeostasis leads to abortive infection outcomes, whereas our findings reveal a complementary viral strategy in which phage portal proteins suppress RelA- and SpoT-mediated (p)ppGpp synthesis during productive infection. Together, these observations suggest that alarmone metabolism constitutes a regulatory interface engaged by both bacterial defense modules and phage countermeasures.

Although our analysis focuses primarily on coliphages, the SR is universal across bacterial phyla, raising the possibility that alarmone signaling may represent a widespread target for viral counter-defense strategies. It remains to be determined whether portal-mediated suppression of (p)ppGpp synthesis extends beyond enterobacterial phages. Moreover, while our biochemical data demonstrate selective inhibition of RelA and SpoT synthetase activity, the precise molecular mechanisms underlying this inhibition - such as whether it involves allosteric regulation or steric occlusion - remain to be resolved. Finally, because (p)ppGpp signaling intersects with multiple defense systems and stress-response pathways, suppression of the SR may indirectly influence other layers of bacterial defense. Dissecting these interactions will be an important avenue for future work aimed at understanding how global physiological defenses integrate into the broader phage**-**host arms race.

## Supplementary figures

**Supplementary Fig. 1.**
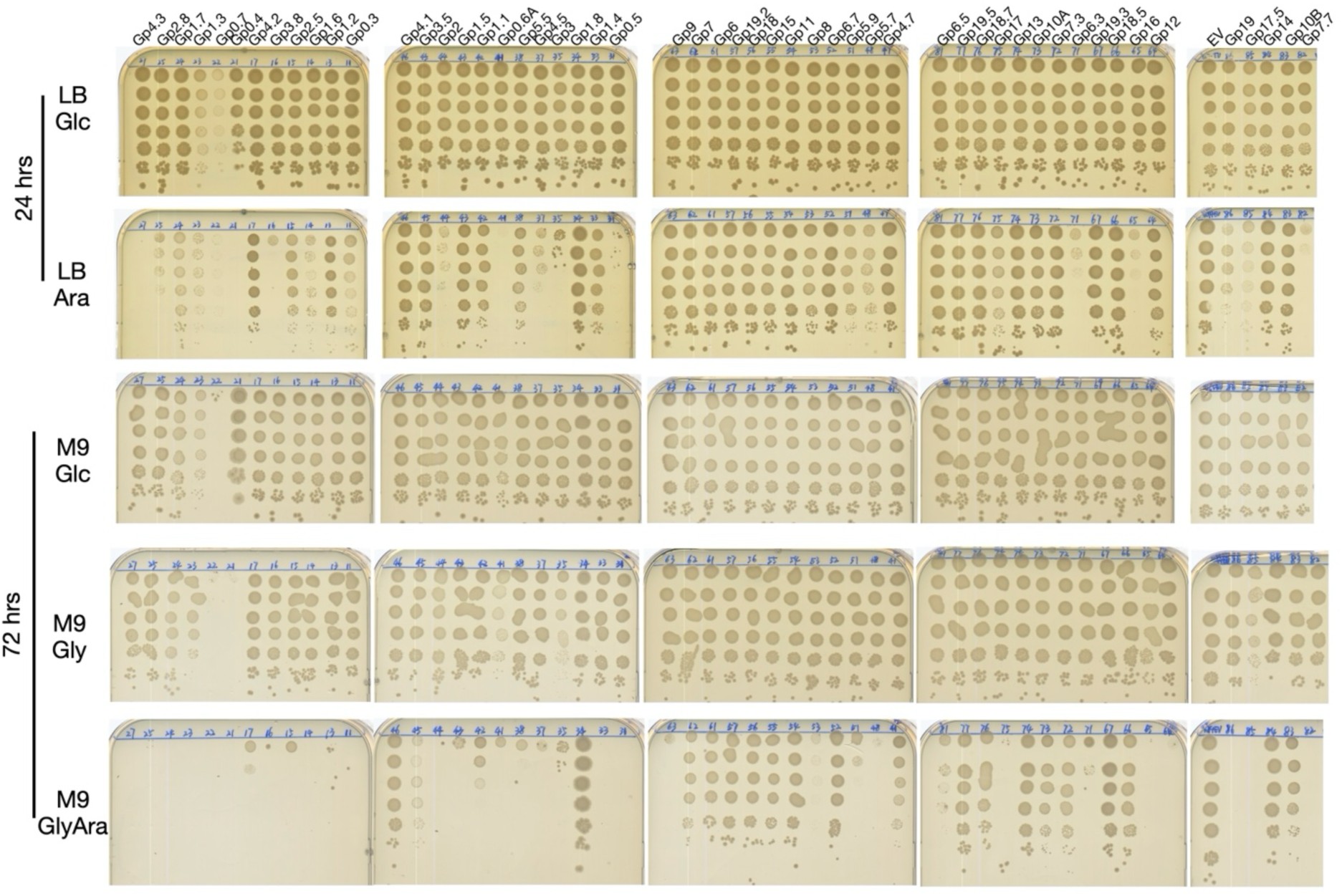
Conditional toxicity of T7 proteins under nutrient limitation. Serial dilution assays showing growth of *Escherichia coli* expressing individual T7 genes in rich and minimal media. Colony-forming units were normalized to OD_600nm_. Expression of multiple T7 proteins resulted in growth inhibition specifically under minimal-medium conditions, whereas little or no effect was observed in rich medium. LB Glc and LB Ara are LB media supplemented with 0.2% glucose and 0.2% arabinose, respectively. M9 Glc, M9 Gly, and M9 Gly Ara denote M9 minimal media supplemented with 0.4% glucose, 0.4% glycerol, or 0.4% glycerol plus 0.2% arabinose, respectively. LB plates were imaged after 24 h, and M9 plates after 72 h.

**Supplementary Fig. 2.**
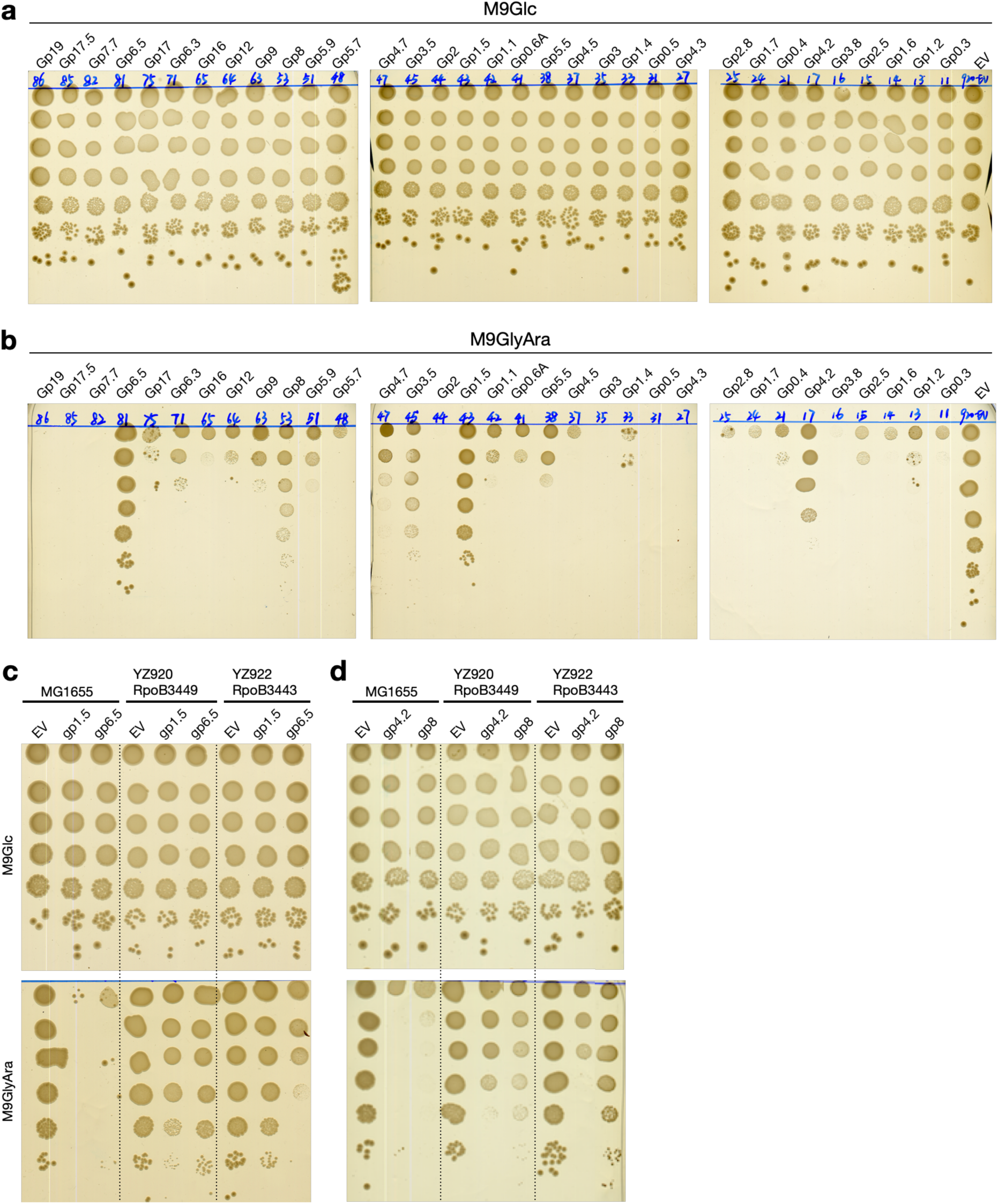
Suppression of T7 protein-induced growth defects by stringent-like RNA polymerase alleles. **a,b**, The YZ920 strain carrying the stringent-like *rpoB*3449 mutation was transformed with pBAD33 constructs expressing T7 proteins that were toxic in the wild-type *E. coli* MG1655 strain. Transformants were tested on M9 medium supplemented with glucose (M9Glc; **a**) or glycerol plus arabinose (M9GlyAra; **b**) and incubated for 72 h. **c,d**, Ten-fold serial dilutions (top to bottom) of wild-type MG1655 or YZ920 and YZ922 strains carrying the stringent-like *rpoB*3449 and *rpoB*3443 mutations, respectively, were spotted onto M9Glc or M9GlyAra plates. Cells expressed the indicated T7 genes (*gp4.2*, *gp1.5*, *gp8*, *gp6.5*) or an empty vector (EV). Plates were incubated at 37 °C and photographed after 72 h.

**Supplementary Fig. 3.**
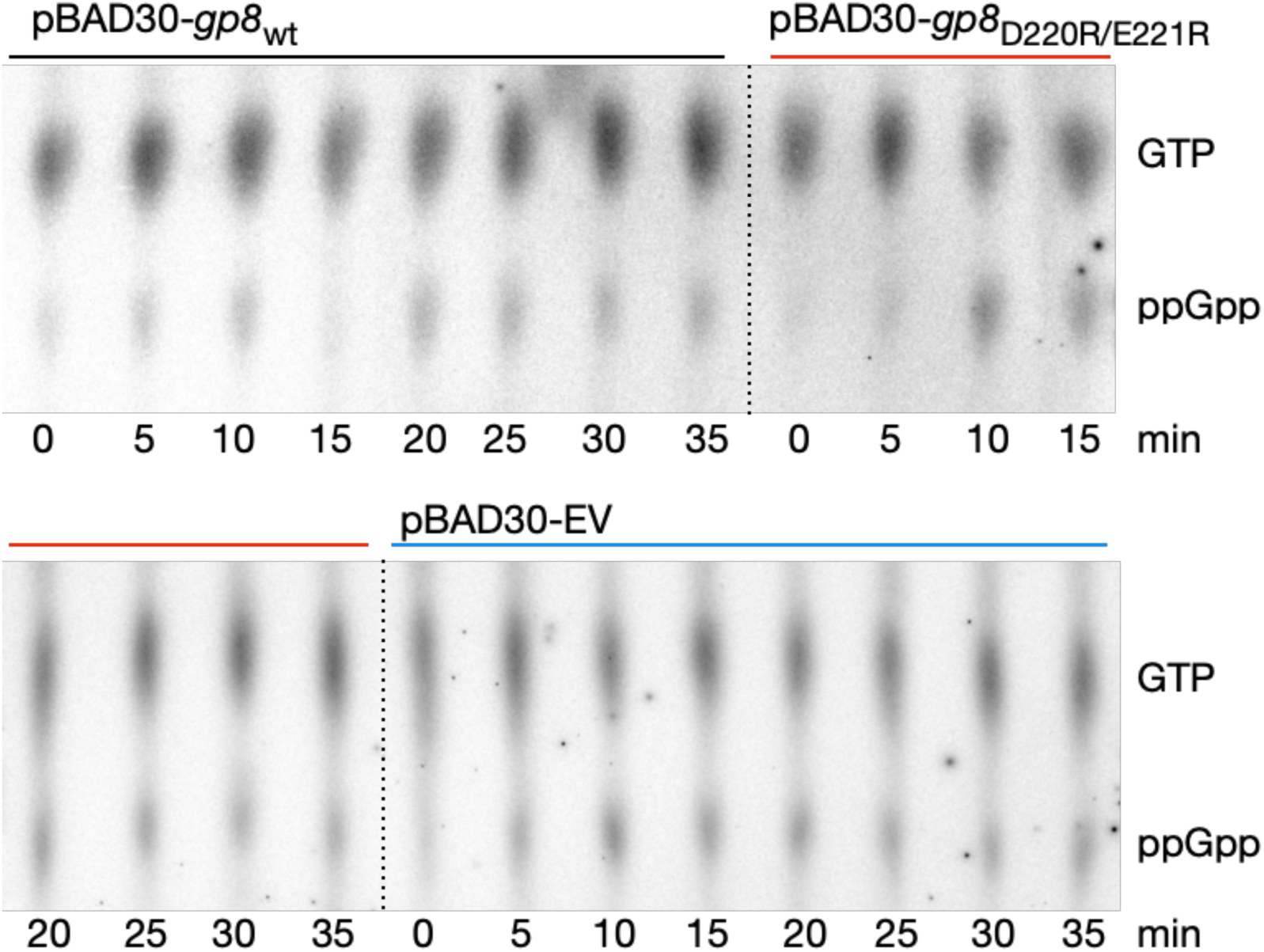
Gp8 expression reduces intracellular (p)ppGpp accumulation. Thin-layer chromatography (TLC) analysis of **³²P**-labeled nucleotide pools during expression of wild-type Gp8 or the interaction-defective mutant Gp8_D220R/E221R_ from the pBAD30 vector. Gp8 expression was induced with 0.2% arabinose, and 25 µL of each sample was analyzed by TLC (see Materials and Methods for details). Positions of GTP and ppGpp are indicated, along with the time points (minutes) following arabinose induction.

**Supplementary Fig. 4.**
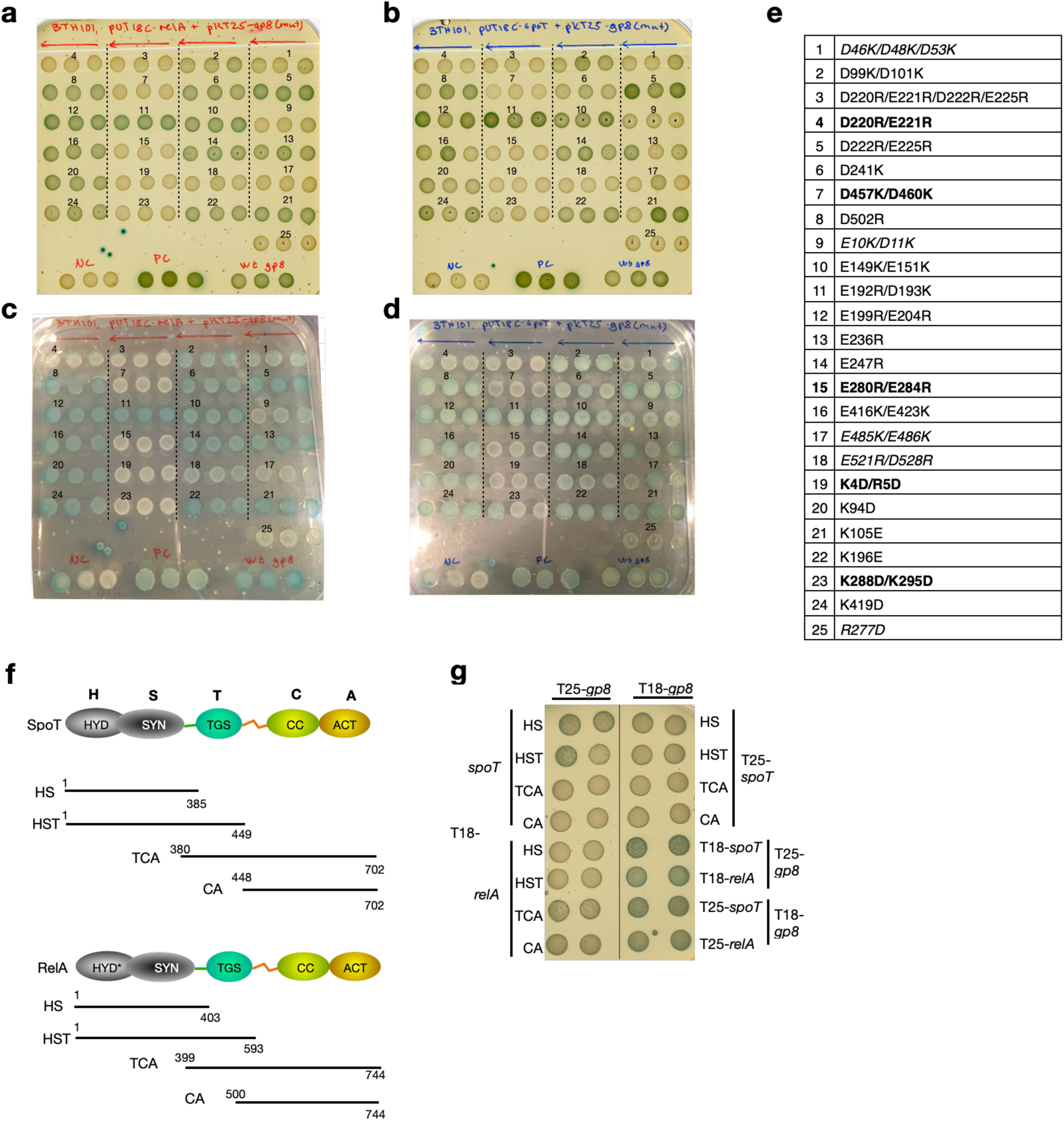
Charge-reversal mutations in Gp8 disrupt interaction with RelA and SpoT. **(a-d)** Bacterial two-hybrid assays assessing interactions between charge-reversal mutants of Gp8 and RelA **(a, c**; 24 h images) or SpoT **(b, d**; 48 h images). Three independent replicate spots are shown for each mutant. Mutations are numbered as indicated in panel **(e)**. Mutants shown in **bold** denote strongly defective interactions, whereas mutants shown in *italics* exhibit partial defects. **(f)**. Schematic overview of RelA and SpoT domain architectures and the truncation constructs tested in panel (**g**). Start and end residue numbers are indicated. (**g**) Bacterial two-hybrid assays assessing interactions between Gp8 and the RelA/SpoT truncations shown in (**f**). Two independent replicates are shown side by side.

**Supplementary Fig. 5.**
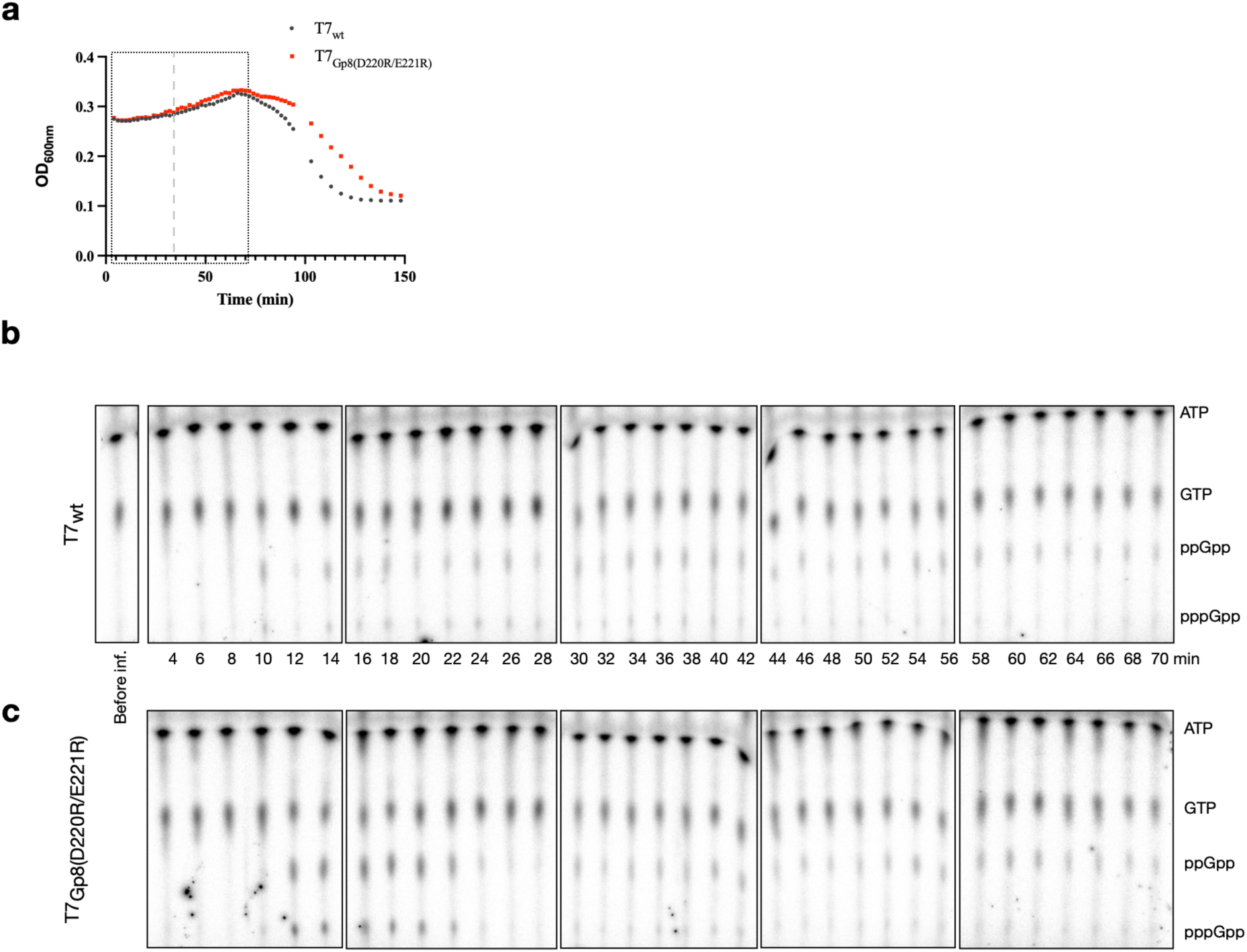
Sustained alarmone accumulation during infection with interaction-defective portal mutants. **(a)** Lysis kinetics of *E. coli* MG1655 growing in in MOPS minimal medium supplemented with low phosphate (0.2 mM K_2_HPO_4_), and 0.2% (v/v) glucose, and infected with wild-type T7 or gp8 interaction-defective mutant phages. The boxed region indicates the time window quantified in **(b,c)** and shown in Fig. 5e. **(b,c)** Radiolabeled nucleotide analysis of alarmone dynamics during phage infection. Intracellular nucleotide pools were ^32^P-labeled during infection of *E. coli* MG1655 in MOPS medium with T7 encoding wild-type gp8 **(b)** or interaction-defective gp8 **(c)**. The positions of ATP, GTP, ppGpp, and pppGpp are indicated.

**Supplementary Fig. 6.**
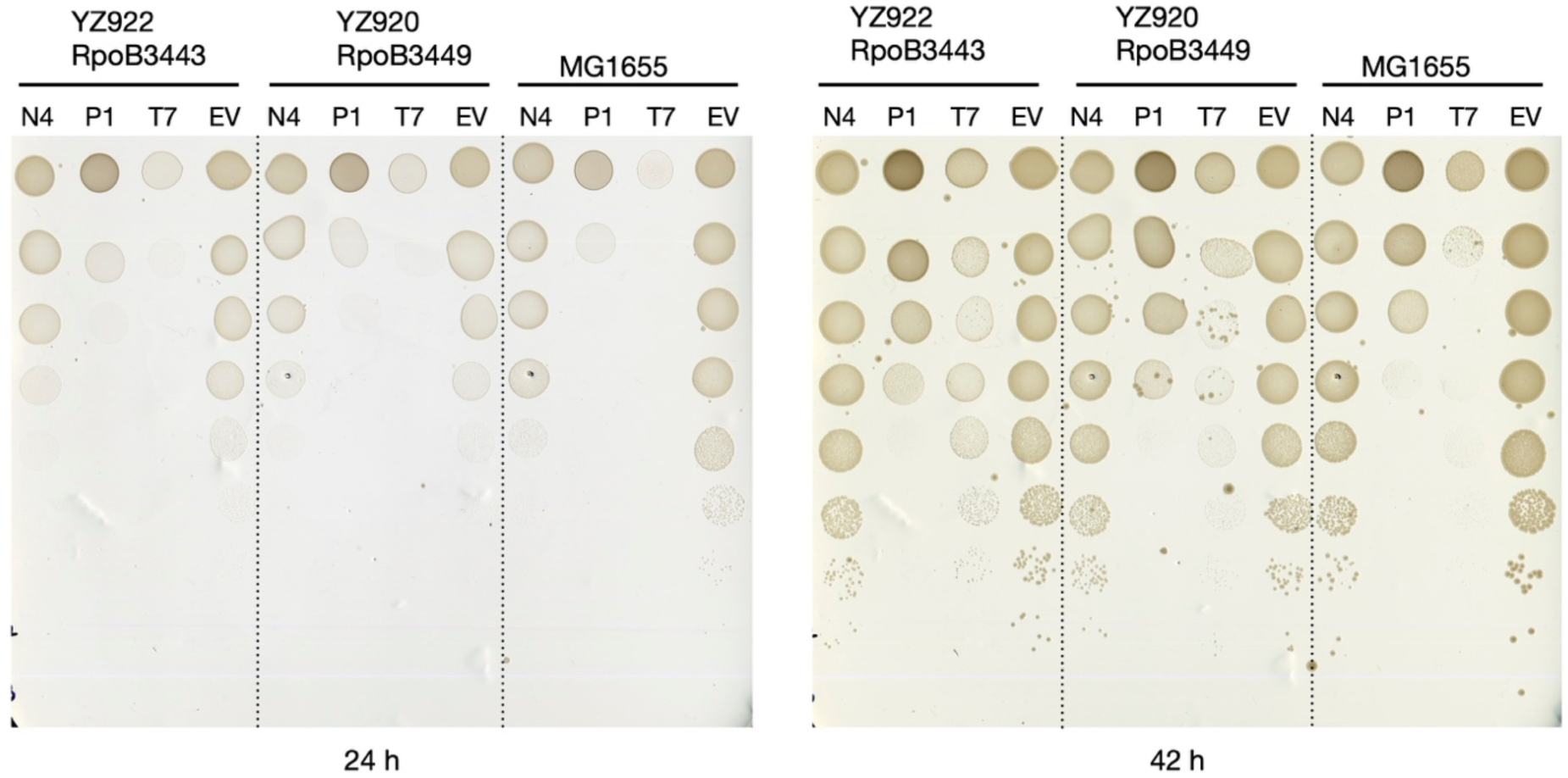
Portal proteins from P1 and N4 exhibit stringent-response-linked phenotypes. Portal proteins from bacteriophages N4 and P1 were cloned into the arabinose-inducible pBAD33 vector and expressed in wild-type *Escherichia coli* MG1655 or in stringent-response**-**altered strains YZ920 and YZ922, carrying the *rpoB3449* and *rpoB3443* alleles, respectively. Toxicity was assessed on M9 minimal medium supplemented with 0.4% glycerol and 0.2% arabinose. Plates were imaged after 24 and 42 h of incubation at 37 °C. Labels N4, P1, and T7 denote portal proteins from the respective phages; EV indicates the empty-vector control.

## Materials and Methods

### Bacterial strains and growth conditions

All *Escherichia coli* strains and primers used in this study are listed in Supplementary Document 1 and 3 respectively. Unless otherwise stated, *E. coli* K-12 MG1655 was used as the wild-type background for infection assays and physiological analyses, and all derivative strains were constructed in this background. *E. coli* DH5α was used for routine cloning, and *E. coli* BL21(DE3) was used for protein expression. Bacteria were routinely grown in lysogeny broth (LB) at 37 °C with aeration, with antibiotics added as required. For nucleotide labeling experiments, cells were grown in defined MOPS minimal medium formulated with low phosphate (0.2 mM K_2_HPO_4_), and 0.2% (v/v) glucose^36^, to minimize background labeling from inorganic phosphate. For experiments in defined nutrient conditions, cells were grown in M9 minimal medium supplemented with glucose (0.2% w/v), glycerol (0.4% v/v) as indicated. Arabinose-inducible expression was achieved by addition of L-arabinose to a final concentration of 0.2% (w/v). Growth and induction conditions are specified in the corresponding figure legends.

### Phage strains and propagation

Phages were propagated using *E. coli* MG1655 as host. A 50 mL MG1655 culture in LB supplemented with 1 mM MgCl_2_ and 1 mM CaCl_2_ was grown at 37°C with shaking at 160 rpm to an OD_600nm_ of ∼ 0.5. Phage stock (100 µL; PFU ∼10^8^-10^10^) was added and incubation was continued until extensive lysis was observed (1-3 h for T7). The lysate was treated with chloroform (1 mL per 10 mL culture), vortexed, incubated for 5-10 min at room temperature, and centrifuged (5 min, 5000 rpm). The supernatant was transferred to a new tube, centrifuged again and collected into a clean tube. The phage stock was concentrated, and the buffer was exchanged following the protocol described here^48^. Briefly, a 5× PEG solution (5 M NaCl, 20%

(w/v) PEG800) was added to the cleared lysate and incubated overnight at 4°C. The precipitated phages were pelleted (1 h, 4°C, 19000 g), resuspended in 0.5-1 mL phage buffer (150 mM NaCl, 40 mM Tris, 10 mM MgSO_4_), and PFU was determined via plaque assay. Phage stocks were stored at 4°C.

#### Liquid killing assay

*E. coli* strains MG1655, *ΔrelA*, and *ΔrelAΔspoT*, were incubated overnight in LB medium at 37°C with shaking at 160 rpm. The following day, cultures were backdiluted into fresh LB medium and incubated in flasks at 37°C with shaking at 160 rpm until reaching an OD_600nm_ of ∼ 0.4. Cultures were normalized to OD_600nm_=0.4, and 150 µL normalized culture was mixed with T7 phage in a clear flat-bottom 96-well plate to the MOI=0.1. The OD_600nm_ was recorded via a plate reader (BioTek) every five minutes at 37°C without shaking.

#### Plaque assay

Plaque assays were performed as described here^49^. Briefly, cultures of the appropriate strains were prepared in LB with appropriate antibiotics and incubated overnight at 37°C shaking at 160 rpm. The following day, cultures were adjusted to OD_600nm_=2, and 200 µL normalized culture was mixed with 100 µL of 10-fold serially diluted phage and incubated at room temperature for 10 min. The mixtures were then mixed with 3 ml melted 0.7% LB agar (kept at 50°C) and immediately poured onto prewarmed 1% LB agar plates. Plates were incubated overnight at 37°C. For time-resolved plaque growth measurements, plates were imaged at 37 °C using the Reshape machine, and plaque diameter expansion over time was quantified using ImageJ.

#### T7 phage engineering

To enable targeted mutagenesis of the essential T7 *gp8* gene, we established a genetic dependency between the host and the phage based on the *trxA* gene, which is a host factor essential for T7 replication. The endogenous *trxA* gene was deleted from the host chromosome, rendering the host non-permissive to wild-type T7 infection. To restore infectivity, *trxA* was introduced into the T7 genome via homologous recombination in place of *gp8*, and the *gp8* was instead provided on the plasmid vector pCA24N^50^ transformed to this *ΔtrxA* host. This created an interdependent system where the *gp8* sequence on the plasmid could be easily manipulated. Due to homology sequences, the plasmid-encoded *gp8* mutant could integrate into its original position on the T7 genome and thus create a mutant T7 particle that can be counter selected using its independence of a plasmid borne *gp8*.

#### Delete trxA in MG1655 using P1 phage transduction

To delete *trxA* from *E. coli* MG1655, a P1 lysate was first prepared using *E. coli ΔtrxA* strain from the Keio collection^51^. Then, the P1 transduction method was used to generate a MG1655 *ΔtrxA::kan* strain^52^ and the kanamycin resistance maker was flipped out by using pCP20^53^.

#### Replace gp8 with trxA in T7

The *trxA* gene was PCR-amplified using genomic DNA from *E. coli* strain MG1655 as a template and primers pYZ1189 and pYZ1190, each containing 40 bp extensions homologous to sequences flanking the *gp8* locus in the T7 phage genome. The purified PCR product was introduced into *E. coli* MG1655 cells by electroporation during co-infection with T7 phage. Immediately following electroporation, LB medium was added, and the culture was incubated at 37°C,160 rpm until lysis was observed (around 1.5 hours). Phage lysates were then harvested with chloroform and clarified via centrifugation (5 min, 5000 rpm).

#### Cloning of pCA24N-gp8 and quick-change mutagenesis of gp8

The *gp8* gene as well as 350 bp flanking regions from T7 was PCR amplified using primers pYZ1191 and pYZ1192 that also encoded a *XhoI* and *HindIII* restriction site, respectively. This PCR product as well as the pCA24N^50^ plasmid were both digested with *XhoI* and *HindIII* and ligated together using T4 ligase. The constructed pCA24N-gp8 was purified and used as template for quick-change mutagenesis with primers pYZ1085-pYZ1136 (see Primer list).

#### Constructing mutant T7 phages with gp8 mutations

pCA24N plasmids encoding mutant *gp8* were purified and transformed into *E. coli* MG1655 *ΔtrxA*. This strain was then infected with T7 *gp8::trxA* phage lysate (prepared as described above). Through homologous recombination between the plasmid-encoded *gp8* mutant and the corresponding flanking regions in the T7 genome (where *trxA* is inserted), the mutant *gp8* sequence can replace *trxA*, and results in T7 progeny that carry the desired *gp8* mutation. The mutant T7 phage particles can be counter selected using its independence of the pCA24N-gp8.

### Measurement of (p)ppGpp by radiolabeling and autoradiography

(p)ppGpp is a very labile molecule not suited for analysis by HPLC or mass spectrometry^54^. We therefore, performed thin layer chromatography (TLC) to quantify both pppGpp and ppGpp. As phosphates in the medium impairs efficient incorporation of H_3_^32^PO_4_ during labelling, MOPS medium supplemented with low phosphate concentration (0.2 mM K_2_HPO_4_), and 0.2% (v/v) glucose was used^36^. MG1655 was grown for ca. 18 hrs in MOPS-lowP_i_-Glc medium at 37 °C with agitation (160 rpm). Afterwards, the strains were adjusted to OD_600nm_=0.05 in MOPS-lowP_i_-Glc and grown at 37 °C with agitation (160 rpm) until OD_600nm_ of 0.2-0-3 was reached. Then, cells were transferred to a 2-ml Eppendorf tube and H_3_^32^PO_4_ (equal to 100 μCi/ml) (Revvity, NEX053001MC) was added. After 1.5 h at 37 °C with 600 rpm agitation, cells were transferred to a new tube and infected with T7 phage to an MOI of 0.1. Immediately after, 150 µl infected cells were transferred to a clear 96-well plate and OD_600nm_ at 37°C was monitored every 2 min via a plate reader (BioTek). Simultaneously, 25 µl sample was collected every 2 min and transferred to 5 µl cold 2 M formic acid. Samples were stores at -20°C until analysed by TLC using 1.5 M KH_2_PO_4_ (pH 3.4) as mobile phase.

### Cloning all T7 genes into the pET28b and pBAD33 vectors

Genomic DNA of *E. coli* T7 bacteriophage was used as the template for amplification of individual T7 genes with Q5 High-Fidelity DNA polymerase (New England Biolabs). PCR amplicons were gel-purified, quantified spectrophotometrically, and digested with *EcoRI* and *NcoI* at 37°C for 3 h followed by enzyme inactivation at 80°C for 20 min. The digested fragments were ligated into the similarly treated pET28b vector using T4 DNA ligase at room temperature for 30 min. Ligation mixtures were transformed into chemically competent *E. coli* DH5α cells and plated on LB agar supplemented with kanamycin (25 µg mL⁻¹). Individual transformants were screened by colony PCR, and clones displaying expected insert sizes were cultured in LB medium for plasmid purification. All constructs were verified by Sanger sequencing, yielding a pET28b**-**T7 gene library encompassing 51 of 59 targeted genes; 8 highly toxic genes could not be cloned or yielded unstable plasmids.

To place the T7 genes under the arabinose-inducible promoter, verified pET**-**T7 plasmids were digested with *XbaI*, *HindIII* and *EcoRV*, and ligated into pBAD33 linearized with *XbaI* and *HindIII*. This procedure transferred the pET Shine-Dalgarno sequence and corresponding T7 gene inserts into pBAD33 under the control of the pBAD promoter. Using the iVEC cloning strategy^55^, we successfully obtained pBAD33-*gp0.4* and pBAD33-*gp0.7* plasmids, the pET versions of which were impossible. The resulting library, designated pBAD**-**T7, comprised 53 T7 genes. For expression of T7 proteins in strain YZ921, the T7 genes were subcloned from the pBAD33 (cat) vector into the pBAD30 (bla) vector using the *XbaI* and *HindIII* restriction sites, as YZ921 carries a chloramphenicol resistance gene (cat).

### Toxicity assay of pBAD33-T7 genes on five media

Each of the 53 verified pBAD**-**T7 plasmids was individually transformed into chemically competent *E. coli* MG1655 cells. Transformants were streaked as single colonies on LB agar plates supplemented with chloramphenicol (25 µg mL⁻¹) and a single colony from each transformant was inoculated into 1 mL LB broth and cultured overnight at 37°C with shaking (200 rpm). From each culture, 300 µL was diluted in 800 µL fresh LB to measure optical density at 600 nm (OD_600nm_), while another 300 µL aliquot was pelleted by centrifugation at 13,000 rpm for 3 min, washed twice with 1 mL PBS, and resuspended to an OD_600nm_ = 1. Serial tenfold dilutions (10⁸**-**10¹ CFU mL⁻¹) were prepared in 96-well plates by mixing 20 µL of bacterial suspension with 180 µL PBS per well. From each dilution, 5 µL was spotted onto five pre-dried chloramphenicol-containing agar media: LB + 0.4% glucose, LB + 0.2% L-arabinose, M9 + 0.4% glucose, M9 + 0.4% glycerol, and M9 + 0.4% glycerol + 0.2% L-arabinose. Plates were incubated at 37°C and photographed every 24 h to monitor colony growth and induction-dependent phenotypic changes.

### Construction of bacterial two-hybrid vectors

#### Cloning relA, spoT and their truncates

*spoT* and *relA* were cloned into the bacterial two-hybrid vectors pUT18C and pKT25. The full-length genes were amplified from *E. coli* MG1655 chromosomal DNA using primer pairs pYZ479/pYZ480 (*spoT*) and pYZ481/pYZ482 (*relA*). PCR products and recipient plasmids were digested with *XbaI* and *XmaI*, ligated, and transformed into *E. coli*. Truncated *spoT* variants corresponding to the HS, HST, TCA, and CA regions were generated by PCR using primer pairs pYZ473/pYZ474, pYZ473/pYZ475, pYZ477/pYZ476, and pYZ478/pYZ476, respectively. The resulting fragments were digested with *XbaI*/*XmaI* and cloned into pUT18C or pKT25. Truncated *relA* variants (HS, HST, TCA, and CA) were amplified using primer pairs pYZ917/pYZ919, pYZ917/pYZ918, pYZ920/pYZ921, and pYZ920/pYZ922, respectively, digested with *XbaI*/*XmaI*, and cloned into pUT18C.

#### Cloning of the portal protein from phages P1 and N4 into pKT25

Portal protein from phages P1 and N4 were amplified using primer pairs pYZ1285/pYZ1286 and pYZ1287/pYZ1288 for phage P1 and N4, respectively. The pKT25 was digested with *KpnI* and *EcoRI*, assembled with both PCR products by the iVEC cloning method^55^.

#### pUT18-rpoZ and pKNT25-rpoZ

To generate pUT18-*rpoZ*, the full-length *rpoZ* gene was PCR-amplified from *E. coli* MG1655 genomic DNA using primers pYZ867 and pYZ868. The amplified fragment and pUT18 vector were digested with *HindIII* and *XbaI*, ligated, and transformed into *E. coli* DH5α. For construction of pKNT25-*rpoZ*, the pKNT25 vector was first modified by site-directed mutagenesis using primers pYZ871 and pYZ872 following the QuickChange protocol to introduce an *NcoI* restriction site, generating pKNT25-*NcoI*. The *rpoZ* insert was excised from pUT18-rpoZ by digestion with *ScaI*, *HindIII*, and *XbaI*, and ligated into *HindIII*/*XbaI*-digested pKNT25-*NcoI*.

#### pUT18/KT25-gp4.2/gp1.5/gp8/gp6.5

An *NcoI* restriction site was introduced into pUT18C (YZ709) by site-directed mutagenesis using primers pYZ864 and pYZ866, generating pUT18C-*NcoI*. For pKT25 (YZ705), the native *NcoI* site was first removed using primers pYZ863 and pYZ862, followed by introduction of a new *NcoI* site using primers pYZ864 and pYZ865 to generate pKT25-*NcoI*. Plasmids pET28-gp4.2, pET28-gp1.5, pET28-gp8, and pET28-gp6.5 were digested with *EcoRI*, *NcoI*, and *HindIII*, and the resulting inserts were individually cloned into pUT18C-*NcoI* or pKT25-*NcoI* vectors digested with *NcoI* and *HindIII*.

#### Construction of Gp8 truncations

Truncations of Gp8 were generated by site-directed mutagenesis of the *gp8* gene on the pKT25 vector in *E. coli* DH5a using the QuickChange method. Primer pairs pYZ1063/pYZ1064, pYZ1065/pYZ1066, pYZ1067/pYZ1068, and pYZ1069/pYZ1070, were used to construct the clip truncation, clip+stem truncation, N-terminal truncation, and C-terminal truncation, respectively. PCR reactions were carried out with Q5 DNA polymerase using an extension time of 3 min and an annealing temperature gradient of 50-70°C across eight 12 µL samples. After amplification, the samples were pooled and digested with 2.5 µL *DpnI* at 37°C for 3 h, followed by inactivation at 80°C for 20 min. The reaction mixture was then transformed into *E. coli* DH5a via chemical transformation. Successful truncations were confirmed by Sanger sequencing using primers pYZ471 and pYZ472.

### Bacterial two-hybrid assay

The bacterial two-hybrid assay was performed using *Escherichia coli* strain BTH101. Cells were co-transformed with plasmids pUT18C and pKT25, with one plasmid encoding either *relA* or *spoT* and the other encoding the phage gene of interest. Transformants were selected on LB agar supplemented with 0.2% (w/v) glucose, 25 µg mL⁻¹ kanamycin, and 100 µg mL⁻¹ ampicillin and incubated at 37 °C. Single colonies were used to inoculate overnight cultures grown at 37 °C with shaking at 170 rpm. The following day, cultures were washed twice with phosphate-buffered saline (PBS) and resuspended in PBS to a final optical density of OD_600nm_ = 5. Aliquots (5 µL) of the normalized cell suspensions were spotted onto LB agar containing 25 µg mL⁻¹ kanamycin, 100 µg mL⁻¹ ampicillin, and 40 µg mL⁻¹ X-gal. Plates were incubated at 30 °C overnight. Plates were scanned, and spot intensities were quantified using ImageJ. Images were converted to RGB stacks, the red channel was isolated and inverted, and the mean intensity of each spot was measured. Background signal from negative controls was subtracted from all samples. Interaction strengths were normalized and expressed as a percentage of the positive control or full-length protein interaction, as specified in the figure legends.

### Cloning of the pET28-HMS-relA and -spoT vectors

#### *pET28-HMS-spoT* and derivatives

Plasmids pET28a and pMAL-c2x were digested with *NdeI* and *EcoRI* and ligated to generate pET28-His₆-MBP (YZ913). The *sumo* and *spoT* genes were PCR-amplified using primers pYZ692/693 and pYZ694/695, fused by overlap PCR, and cloned into pET28a using *NcoI* and *HindIII* to generate pET28a-His₆-SUMO-SpoT (YZ307). The SUMO-SpoT fragment was excised with *EcoRI* and *HindIII* and ligated into pET28a-His₆-MBP. Missing SUMO sequences were amplified using primers pYZ684/685, digested with *EcoRI*, and inserted into this intermediate construct to generate pET28-His₆-MBP-SUMO-SpoT (pET28-HMS-spoT).

#### Generation of SpoT point mutants

The E319Q and H72A/D73A mutations were introduced into pET28-HMS-spoT using PCR-amplified fragments derived from chromosomal *spoT* mutant strains YZ586 and YZ905, respectively^56^. PCR products generated with primers pYZ702 and pYZ703 were digested with *PmlI* and *SalI* and ligated into pET28-HMS-spoT to generate the corresponding mutant constructs. The *spoT* gene was deleted from pET28-HMS-spoT using primers pYZ905 and pYZ906 to generate the pET28b-HMS vector (YZ1515).

#### Construction of *pET28-HMS-relA*

The *relA* gene was PCR-amplified from *E. coli* MG1655 using primers pYZ927 and pYZ928. Both the PCR product and pET28b-HMS were digested with *BamHI* and *HindIII*, ligated using T4 DNA ligase, and transformed into *E. coli* DH5α to generate pET28-HMS-relA.

### Pull down and Western blot

Overnight cultures of *E. coli* BL21 DE3 strains expressing pRARE, pET28b-HMS, -*relA* or - *spoT*, and pBAD30-*gp8-Flag* (see Strain list) were incubated in LB medium with 25 µg/mL kanamycin, 25 µg/mL chloramphenicol, and 30 µg/mL ampicillin at 37°C with shaking at 160 rpm. The following day, cultures were back-diluted to an OD_600nm_ of 0.05 in 50 mL fresh LB medium containing the same antibiotics as above and incubated at 37°C with shaking at 160 rpm until reaching OD_600nm_ of ∼ 0.2-0.3. Gene expression was induced by addition of arabinose and IPTG to final concentrations of 0.2% and 0.5 mM, respectively, followed by incubation for 4 hours at 37°C with shaking at 160 rpm. Cells were harvested by centrifugation (10 min, 4000 rpm, 4°C), resuspended in 10 mL PBS and centrifuged again as above. Supernatant was discarded and cell pellets were stored at -20°C. Pellets were thawed on ice, resuspended in 400 µL lysis buffer (50 mM HEPES (pH 8), 0.25 M NaCl, 10 mM imidazole, 200 µM ZnCl_2_, and 2 mM MgCl_2_) and disrupted with glass beads (Sigma, G4649) using a MagNA Lyser (20 s at 6500 rpm, done twice with cooling on ice between runs). Lysates were centrifuged (5 min, 5000 rpm, 4°C) and supernatants were collected. Total protein concentration was measured via Bradford Assay, and samples were normalized with lysis buffer to equal total protein concentrations. Equal volumes of normalized lysates were incubated with 50 µL Ni-NTA Agarose resins (Qiagen) overnight at 4°C with gentle inversion. The next day, samples were centrifuged (2 min, 1000 rpm, 4°C), and supernatant was removed. Resins were gently resuspended with 1 mL lysis buffer and incubated for 1 h at 4°C with gentle inversion. Samples were centrifuged (2 min, 1000 rpm, 4°C), and supernatant was removed. Resins were subsequently resuspended in 1 mL wash buffer (50 mM HEPES (pH 8), 0.25 M NaCl, 20 mM imidazole, 200 µM ZnCl_2_, 2 mM MgCl_2_) and incubated for 1 h at 4°C with gentle inversion. Following incubation, samples were transferred to a 0.5 mL Eppendorf tube and centrifuged (first for 2 min at 1000 rpm, then for 1 min at 13000 rpm). Supernatant was removed and proteins were eluted with 20 µL elution buffer (50 mM HEPES (pH 8), 0.25 M NaCl, 1.25 M imidazole, 200 µM ZnCl_2_, 2 mM MgCl_2_) to yield a final imidazole concentration of approximately 500 mM). Samples were incubated for 15 min at 4°C (gently disturbing the resins occasionally). Samples were then centrifuged (2 min, 1000 rpm), and 35 µL supernatant was transferred to 11.6 µL 4x Leammli buffer, boiled for 15 min at 95°C and 12 µL of each sample was loaded onto a 10% SDS-PAGE gel. Following electrophoresis, proteins were transferred to a PVDF membrane by wet transfer (300 mA, 1 h, 4°C) using transfer buffer composed of 20% methanol, 0.2 M glycine, and 25 mM Tris (pH 8.3). Membrane was blocked overnight at 4°C in TBST + 5% (w/v) milk powder. Next day, the membrane was incubated with anti-FLAG primary antibody diluted in TBST + 5% (w/v) milk powder for 1 h, 4°C, washed three times (10 min each) in TBST, and subsequently incubated with anti-mouse secondary antibody in TBST + 5% (w/v) milk powder for 1 h, 4°C. After three additional washes in TBST, the membrane was developed using Pierce^®^ ECL Western Blotting Substrate (Thermo Scientific, #32209) and imaged via ImageQuant LAS4000.

### Protein purification for biochemical assays

RelA (YZ1607), SpoT hydrolysis only (HYD, YZ1063), SpoT synthesis only (SYN, YZ1070), and Gp8 (YZ1632) constructs bearing an N-terminal His**-**MBP**-**SUMO (HMS) tag were expressed from pET28b in *E. coli* BL21(DE3). Overnight cultures of these strains were grown in LB medium supplemented with appropriate antibiotics at 37°C, with shaking at 160 rpm. The following day, cultures were back-diluted 1:50 into 0.5 L fresh LB medium supplemented with appropriate antibiotics and grown at 37°C with shaking at 160 rpm until reaching an OD of 0.4-0.7. Protein expression was induced by the addition of IPTG to a final concentration of 0.1 mM, and cultures were incubated overnight at 24°C, shaking with aeration. The next day, cells were harvested by centrifugation (5000 rpm, 10 min, 4°C), washed in 30 mL cold PBS, and pelleted again (4000 rpm, 20 min, 4°C). Resulting pellets were resuspended in 40 mL cold lysis buffer (50 mM Tris-HCl pH=8, 1 M NaCl, 2 mM DTT, 10 mM imidazole) supplemented with Pierce^TM^ Protease Inhibitor Mini (Thermo Scientific, A32955). Cells were lysed by sonication on ice for 24 min at 20% amplitude, and lysates were cleared by centrifugation (14000 rpm for 45 min at 4 °C). Cleared lysates were incubated with 0.5-1 mL Ni-NTA Agarose resins (Qiagen) overnight at 4°C with gentle inversion. The next day, the resins were collected by centrifugation (1000 rpm, 2 min) and washed once with 30 mL lysis buffer (1 h, 4°C), followed by 30 mL wash buffer (same composition as lysis buffer, except imidazole increased to 20 mM). Resins were transferred onto a Bio-Rad Gravity Flow Column, and bound proteins were eluted with 700 µL elution buffer (50 mM tris-HCl pH=8, 1 M NaCl, 2 mM DTT, 500 mM imidazole). The eluted fraction was subjected to size-exclusion chromatography using an ӒKTA Pure^TM^ system (Cytiva) equilibrated gel filtration buffer (50 mM HEPES pH=8, 1 M NaCl, 2 mM DTT, 5% glycerol). Protein concentrations were determined via Bradford Assay.

### Synthesis of [α-³²P](p)ppGpp via RelSeq

[α-³²P](p)ppGpp was synthesized using[α-³²P]GTP (revvity, BLU006X250UC) as substrate in a RelSeq-catalyzed reaction^57^. RelSeq (YZ8) was expressed and purified as described above except that protein expression was induced with 0.5 mM IPTG for 3 h at 30°C. The purification buffer contained 500 mM NaCl, 50 mM Tris-HCl (pH 8), and 5% glycerol, with imidazole concentrations of 20, 40, and 500 mM for lysis, wash, and elution buffers, respectively. The gel filtration buffer was identical but without imidazole. For the enzyme reaction, a 10× reaction buffer was prepared containing 250 mM Tris (pH 9), 1 M NaCl, and 150 mM MgCl_2_. The final reaction system contained 1× reaction buffer, 8 mM ATP, 125 nM [α-³²P]GTP, and 4 µM RelSeq, and was incubated at 37°C for 1 h. The reaction was terminated by boiling for 5 min at 95°C, followed by centrifugation (14000 rpm, 5 min). Conversion of substrate was verified by thin-layer chromatography (TLC) using 0.5 µL of the reaction mixture and 1.5 M KH_2_PO_4_ (pH 3.4) as the mobile phase.

### Biochemical assays

Synthetase activity of RelA and SpoT (SYN), was assayed in reaction mixtures containing 50 mM HEPES (pH 7.4), 100 mM KCl, 5 mM MgCl_2_, 2 mM DTT, 0.2 mg/mL BSA, 2 mM GTP, and 6 nM [α-³²P] GTP (revvity, BLU006X250UC) were prepared. Protein was added to final concentrations of 0.5 µM (RelA) or 3 µM (SpoT (SYN)). Gp8 was added to varying final concentrations, and samples were incubated at room temperature for 10 min before enzyme reactions were initiated by addition of 5 mM ATP. Hydrolase activity of SpoT (HYD) was assayed in a reaction mixture containing 50 mM HEPES (pH 8), 100 mM KCl, 5 mM MnCl_2_, 2 mM DTT, and 0.2mg/mL BSA was prepared. SpoT (HYD) was added to a final concentration of 100 nM, while Gp8 was added at varying final concentrations. Reaction mixtures were incubated at room temperature for 10 min before enzyme reaction was initiated by the simultaneous addition of cold ppGpp (Jena Biocience, NU-884L), and hot [α-³²P](p)ppGpp to final concentrations of 31.25 and 6 nM, respectively. Enzyme reactions were stopped by transferring 4 µL of the reaction mixtures into 1 µL cold 250 mM EDTA (final concentration 50 mM). Samples were loaded onto a PEI Cellulose plates and resolved using 1.5 M KH_2_PO_4_ (pH 3.4) as the mobile phase. Plates were exposed to a BAS-MS imaging plate and scanned using a Typhoon phosphorimager.

## Acknowledgements

We would like to thank Jeppe Aaby Andersen for cloning of the portal protein from phages P1 and N4 into the bacterial two-hybrid system, and for testing their interaction with RelA and SpoT. We would also like to thank Sonia de Araujo and Anette Vestergaard for cloning of the *relA*, *spoT* and their truncates into the bacterial two-hybrid system. We thank Yueyue Liu for preparing stocks of the selected phages obtained from the BASEL Collection. We are grateful for Michael J Gray’s gifting of the RNAP^SR^ strains. The work is supported by a Danmarks Frie Forskningsfond grant (2032-00030B) and the Carlsberg Foundation Semper Ardens Accelerate grant (CF24-1843) to Y.E.Z. and the Chinese Scholarship Council program (202006330098) to L.K.W.

## References

1 Egido, J. E., Costa, A. R., Aparicio-Maldonado, C., Haas, P. J. & Brouns, S. J. J. Mechanisms and clinical importance of bacteriophage resistance. FEMS Microbiol Rev 46 (2022). 10.1093/femsre/fuab048

2 Georjon, H. & Bernheim, A. The highly diverse antiphage defence systems of bacteria. Nat Rev Microbiol 21, 686–700 (2023). 10.1038/s41579-023-00934-x

3 Hochhauser, D. & Sorek, R. Manipulation of the nucleotide pool in human, bacterial and plant immunity. Nat Rev Immunol 26, 7–22 (2026). 10.1038/s41577-025-01206-w

4 Bernheim, A. & Sorek, R. The pan-immune system of bacteria: antiviral defence as a community resource. Nat Rev Microbiol 18, 113–119 (2020). 10.1038/s41579-019-0278-2

5 Lopatina, A., Tal, N. & Sorek, R. Abortive Infection: Bacterial Suicide as an Antiviral Immune Strategy. Annu Rev Virol 7, 371–384 (2020). 10.1146/annurev-virology-011620-040628

6 Potrykus, K. & Cashel, M. (p)ppGpp: still magical? Annual review of microbiology 62, 35–51 (2008). 10.1146/annurev.micro.62.081307.162903

7 Irving, S. E., Choudhury, N. R. & Corrigan, R. M. The stringent response and physiological roles of (pp)pGpp in bacteria. Nat Rev Microbiol 19, 256–271 (2021). 10.1038/s41579-020-00470-y

8 Hauryliuk, V., Atkinson, G. C., Murakami, K. S., Tenson, T. & Gerdes, K. Recent functional insights into the role of (p)ppGpp in bacterial physiology. Nat Rev Microbiol 13, 298–309 (2015). 10.1038/nrmicro3448

9 Nowicki, D., Kobiela, W., Wegrzyn, A., Wegrzyn, G. & Szalewska-Palasz, A. ppGpp-dependent negative control of DNA replication of Shiga toxin-converting bacteriophages in Escherichia coli. J Bacteriol 195, 5007–5015 (2013). 10.1128/JB.00592-13

10 Bryan, D., El-Shibiny, A., Hobbs, Z., Porter, J. & Kutter, E. M. Bacteriophage T4 Infection of Stationary Phase E. coli: Life after Log from a Phage Perspective. Front Microbiol 7, 1391 (2016). 10.3389/fmicb.2016.01391

11 Tabib-Salazar, A. et al. T7 phage factor required for managing RpoS in Escherichia coli. Proc Natl Acad Sci U S A 115, E5353–E5362 (2018). 10.1073/pnas.1800429115

12 Zhang, T. et al. Direct activation of a bacterial innate immune system by a viral capsid protein. Nature 612, 132–140 (2022). 10.1038/s41586-022-05444-z

13 Jimmy, S. et al. A widespread toxin-antitoxin system exploiting growth control via alarmone signaling. Proc Natl Acad Sci U S A 117, 10500–10510 (2020). 10.1073/pnas.1916617117

14 Kurata, T. et al. RelA-SpoT Homolog toxins pyrophosphorylate the CCA end of tRNA to inhibit protein synthesis. Mol Cell 81, 3160–3170 e3169 (2021). 10.1016/j.molcel.2021.06.005

15 Zhang, T. et al. A bacterial immunity protein directly senses two disparate phage proteins. Nature 635, 728–735 (2024). 10.1038/s41586-024-08039-y

16 Ho, P. et al. Bacteriophage antidefense genes that neutralize TIR and STING immune responses. Cell Rep 42, 112305 (2023). 10.1016/j.celrep.2023.112305

17 Niault, T., van Houte, S., Westra, E. & Swarts, D. C. Evolution and ecology of anti-defence systems in phages and plasmids. Curr Biol 35, R32–R44 (2025). 10.1016/j.cub.2024.11.033

18 Murtazalieva, K., Mu, A., Petrovskaya, A. & Finn, R. D. The growing repertoire of phage anti-defence systems. Trends Microbiol 32, 1212–1228 (2024). 10.1016/j.tim.2024.05.005

19 Haseltine, W. A. & Block, R. Synthesis of guanosine tetra- and pentaphosphate requires the presence of a codon-specific, uncharged transfer ribonucleic acid in the acceptor site of ribosomes. Proc Natl Acad Sci U S A 70, 1564–1568 (1973). 10.1073/pnas.70.5.1564

20 Xiao, H. et al. Residual guanosine 3’,5’-bispyrophosphate synthetic activity of relA null mutants can be eliminated by spoT null mutations. J Biol Chem 266, 5980–5990 (1991).

21 Fernández-Coll, L. & Cashel, M. Possible Roles for Basal Levels of (p)ppGpp: Growth Efficiency Vs. Surviving Stress. Front Microbiol 11, 592718 (2020). 10.3389/fmicb.2020.592718

22 Vinella, D., Albrecht, C., Cashel, M. & D’Ari, R. Iron limitation induces SpoT-dependent accumulation of ppGpp in Escherichia coli. Mol Microbiol 56, 958–970 (2005). 10.1111/j.1365-2958.2005.04601.x

23 Spira, B., Silberstein, N. & Yagil, E. Guanosine 3’,5’-bispyrophosphate (ppGpp) synthesis in cells of Escherichia coli starved for Pi. J Bacteriol 177, 4053–4058 (1995). 10.1128/jb.177.14.4053-4058.1995

24 Ross, W., Vrentas, C. E., Sanchez-Vazquez, P., Gaal, T. & Gourse, R. L. The magic spot: a ppGpp binding site on E. coli RNA polymerase responsible for regulation of transcription initiation. Mol Cell 50, 420–429 (2013). 10.1016/j.molcel.2013.03.021

25 Ross, W. et al. ppGpp Binding to a Site at the RNAP-DksA Interface Accounts for Its Dramatic Effects on Transcription Initiation during the Stringent Response. Mol Cell 62, 811–823 (2016). 10.1016/j.molcel.2016.04.029

26 Sanchez-Vazquez, P., Dewey, C. N., Kitten, N., Ross, W. & Gourse, R. L. Genome-wide effects on Escherichia coli transcription from ppGpp binding to its two sites on RNA polymerase. Proceedings of the National Academy of Sciences 116, 8310–8319 (2019). doi:10.1073/pnas.1819682116

27 Durfee, T., Hansen, A. M., Zhi, H., Blattner, F. R. & Jin, D. J. Transcription profiling of the stringent response in Escherichia coli. J Bacteriol 190, 1084–1096 (2008). 10.1128/JB.01092-07

28 Traxler, M. F. et al. The global, ppGpp-mediated stringent response to amino acid starvation in Escherichia coli. Molecular microbiology 68, 1128–1148 (2008). 10.1111/j.1365-2958.2008.06229.x

29 Bae, B. et al. Phage T7 Gp2 inhibition of Escherichia coli RNA polymerase involves misappropriation of sigma70 domain 1.1. Proceedings of the National Academy of Sciences 110, 19772–19777 (2013). doi:10.1073/pnas.1314576110

30 Severinova, E. & Severinov, K. Localization of the Escherichia coli RNA polymerase beta’ subunit residue phosphorylated by bacteriophage T7 kinase Gp0.7. J Bacteriol 188, 3470–3476 (2006). 10.1128/jb.188.10.3470-3476.2006

31 Savalia, D., Robins, W., Nechaev, S., Molineux, I. & Severinov, K. The role of the T7 Gp2 inhibitor of host RNA polymerase in phage development. J Mol Biol 402, 118–126 (2010). 10.1016/j.jmb.2010.07.012

32 Sarubbi, E., Rudd, K. E. & Cashel, M. Basal ppGpp level adjustment shown by new spoT mutants affect steady state growth rates and rrnA ribosomal promoter regulation in Escherichia coli. Molecular and General Genetics MGG 213, 214–222 (1988). 10.1007/BF00339584

33 Grucela, P. K. & Zhang, Y. E. Basal level of ppGpp coordinates Escherichia coli cell heterogeneity and ampicillin resistance and persistence. Microb Cell 10, 248–260 (2023). 10.15698/mic2023.11.808

34 Gray Michael, J. Inorganic Polyphosphate Accumulation in Escherichia coli Is Regulated by DksA but Not by (p)ppGpp. Journal of Bacteriology 201, 10.1128/jb.00664-00618 (2019). 10.1128/jb.00664-18

35 Gray Michael, J. Interactions between DksA and Stress-Responsive Alternative Sigma Factors Control Inorganic Polyphosphate Accumulation in Escherichia coli. Journal of Bacteriology 202, 10.1128/jb.00133-00120 (2020). 10.1128/jb.00133-20

36 Zhang, Y. E. et al. (p)ppGpp Regulates a Bacterial Nucleosidase by an Allosteric Two-Domain Switch. Mol Cell 74, 1239–1249.e1234 (2019). 10.1016/j.molcel.2019.03.035

37 Grucela, P. K., Fuhrer, T., Sauer, U., Chao, Y. & Zhang, Y. E. Ribose 5-phosphate: the key metabolite bridging the metabolisms of nucleotides and amino acids during stringent response in Escherichia coli? Microb Cell 10, 141–144 (2023). 10.15698/mic2023.07.799

38 Wang, B. et al. Affinity-based capture and identification of protein effectors of the growth regulator ppGpp. Nat Chem Biol 15, 141–150 (2019). 10.1038/s41589-018-0183-4

39 Wang, B., Grant, R. A. & Laub, M. T. ppGpp Coordinates Nucleotide and Amino-Acid Synthesis in E. coli During Starvation. Mol Cell 80, 29–42.e10 (2020). 10.1016/j.molcel.2020.08.005

40 Maffei, E. et al. Systematic exploration of Escherichia coli phage-host interactions with the BASEL phage collection. PLoS Biol 19, e3001424 (2021). 10.1371/journal.pbio.3001424

41 Qimron, U., Kulczyk, A. W., Hamdan, S. M., Tabor, S. & Richardson, C. C. Inadequate inhibition of host RNA polymerase restricts T7 bacteriophage growth on hosts overexpressing udk. Mol Microbiol 67, 448–457 (2008). 10.1111/j.1365-2958.2007.06058.x

42 Kingston, R. E., Nierman, W. C. & Chamberlin, M. J. A direct effect of guanosine tetraphosphate on pausing of Escherichia coli RNA polymerase during RNA chain elongation. Journal of Biological Chemistry 256, 2787–2797 (1981). 10.1016/S0021-9258(19)69683-3

43 Tabib-Salazar, A. & Wigneshweraraj, S. RNA Management During T7 Infection. Phage (New Rochelle) 3, 136–140 (2022). 10.1089/phage.2022.0029

44 Rodnina, M. V. Translation in Prokaryotes. Cold Spring Harb Perspect Biol 10 (2018). 10.1101/cshperspect.a032664

45 Casjens, S. R. The DNA-packaging nanomotor of tailed bacteriophages. Nat Rev Microbiol 9, 647–657 (2011). 10.1038/nrmicro2632

46 Stano, N. M. & Patel, S. S. T7 lysozyme represses T7 RNA polymerase transcription by destabilizing the open complex during initiation. J Biol Chem 279, 16136–16143 (2004). 10.1074/jbc.M400139200

47 Zhang, X. & Studier, F. W. Multiple Roles of T7 RNA Polymerase and T7 Lysozyme During Bacteriophage T7 Infection. Journal of Molecular Biology 340, 707–730 (2004). 10.1016/j.jmb.2004.05.006

48 Borges, A. & Athukoralage, J. Phage amplification and concentration, <https://www.protocols.io/view/phage-amplification-and-concentration-yxmvmnb86g3p/v1/references> (2022).

49 Ceelen, M. Plaque assay, <https://www.protocols.io/view/plaque-assay-kqdg35ebqv25/v1> (2019).

50 Kitagawa, M. et al. Complete set of ORF clones of Escherichia coli ASKA library (A Complete Set of E. coli K-12 ORF Archive): Unique Resources for Biological Research. DNA Research 12, 291–299 (2006). 10.1093/dnares/dsi012

51 Baba, T. et al. Construction of Escherichia coli K-12 in-frame, single-gene knockout mutants: the Keio collection. Mol Syst Biol 2, 2006.0008 (2006). 10.1038/msb4100050

52 Thomason, L. C., Costantino, N. & Court, D. L. E. coli genome manipulation by P1 transduction. Curr Protoc Mol Biol Chapter 1, 1 17 11–11 17 18 (2007). 10.1002/0471142727.mb0117s79

53 Cherepanov, P. P. & Wackernagel, W. Gene disruption in Escherichia coli: TcR and KmR cassettes with the option of Flp-catalyzed excision of the antibiotic-resistance determinant. Gene 158, 9–14 (1995). 10.1016/0378-1119(95)00193-a

54 Varik, V., Oliveira, S. R. A., Hauryliuk, V. & Tenson, T. HPLC-based quantification of bacterial housekeeping nucleotides and alarmone messengers ppGpp and pppGpp. Sci Rep 7, 11022 (2017). 10.1038/s41598-017-10988-6

55 Nozaki, S. & Niki, H. Exonuclease III (XthA) Enforces In Vivo DNA Cloning of Escherichia coli To Create Cohesive Ends. J Bacteriol 201 (2019). 10.1128/jb.00660-18

56 Liu, Y. et al. Basal ppGpp regulation by SpoT coordinates metabolic homeostasis and acid resistance. bioRxiv, 2026.2001.2026.700336 (2026). 10.64898/2026.01.26.700336

57 Zhang, Y., Zborníková, E., Rejman, D. & Gerdes, K. Novel (p)ppGpp Binding and Metabolizing Proteins of Escherichia coli. mBio 9 (2018). 10.1128/mBio.02188-17

