## Supplemented document 1 for "Phage portal proteins counteract stringent-response–mediated restriction"

**Supplemented document 1: all strains used in this study**

| **Number** | **Host** | **Plasmid or genotype** | **Reference** |
| --- | --- | --- | --- |
| YZ8 | BL21 DE3 | pENH385-relseq.his | ^1^ |
| YZ15 | DH5a | pCA24N | ^2^ |
| YZ38 | MG1655 | *∆relA* | Laboratory strain |
| YZ44 | BT340 | pCP20 | Laboratory strain |
| YZ142 | MG1655 | Wild-type strain | Laboratory strain |
| YZ213 | DH5a | pBAD30 | Laboratory strain |
| YZ307 | DH5α | pET28-His6.SUMO.spoT |  |
| YZ584 | MG1655 ∆*relA* | *spoT203* (R140C) | ^3^ |
| YZ585 | MG1655 ∆*relA* | *spoT202* (T78I) (4× basal levels of (p)ppGpp) | ^3^ |
| YZ586 | MG1655 ∆*relA* | spoT (E319Q) | ^4^ |
| YZ666 | BTH101 | F-, *cya*-99, *araD*139, *galE*15, *galK*16, *rpsL*1 (Str^R^), *hsdR*2, *mcrA*1, *mcrB*1 | Laboratory strain |
| YZ705 | DH5α | pKT25 | Laboratory strain |
| YZ709 | DH5α | pUT18C | Laboratory strain |
| YZ710 | DH5a | pUT18C-*relA* | This study |
| YZ711 | DH5a | pUT18C-*spoT* | This study |
| YZ712 | DH5a | pKT25-*relA* | This study |
| YZ716 | DH5a | pKT25-*spoT* | This study |
| YZ749 | DH5a | pUT18C-*spoT*(HS) | This study |
| YZ750 | DH5a | pUT18C-*spoT*(HST) | This study |
| YZ751 | DH5a | pUT18C-*spoT*(TCA) | This study |
| YZ752 | DH5a | pUT18C-*spoT*(CA) | This study |
| YZ753 | DH5a | pKT25-*spoT*(HS) | This study |
| YZ754 | DH5a | pKT25-*spoT*(HST) | This study |
| YZ755 | DH5a | pKT25-*spoT*(TCA) | This study |
| YZ756 | DH5a | pKT25-*spoT*(CA) | This study |
| YZ840 | MG1655 | *∆relA∆spoT* | ^3^ |
| YZ905 | MG1655 ∆*relA* | spoT (H72AD73A) | ^4^ |
| YZ913 | DH5α | pET28a-MBP | This study |
| YZ920 | MG1655 | *rpoB3449* (encoding RpoBΔAla532) | Gifted by Michael J Gray^5^ |
| YZ921 | MG1655 *ΔrelA782 spoT207::cat* | *rpoB3449* | Gifted by Michael J Gray^5^ |
| YZ922 | MG1655 | *rpoB3443* | Gifted by Michael J Gray^6^ |
| YZ1063 | Rosetta | pET28-his6.MBP.SUMO.*spoT* (E319Q) | This study |
| YZ1070 | Rosetta | pET28-his6.MBP.SUMO.*spoT* (H72D73AA) | This study |
| YZ1607 | BL21 DE3 | pET28b-his6.MBP.SUMO-*relA* | This study |
| YZ1381 | DH5a | pUT18-*rpoZ* | This study |
| YZ1382 | DH5a | pKNT25-*rpoZ* | This study |
| YZ1383 | DH5a | pUT18C-T7*gp4.2* | This study |
| YZ1385 | DH5a | pUT18C-T7*gp1.5* | This study |
| YZ1387 | DH5a | pUT18C-T7*gp8* | This study |
| YZ1388 | DH5a | pUT18C-T7*gp6.5* | This study |
| YZ1389 | DH5a | pKT25-T7*gp4.2* | This study |
| YZ1391 | DH5a | pKT25-T7*gp1.5* | This study |
| YZ1393 | DH5a | pKT25-T7*gp8* | This study |
| YZ1394 | DH5a | pKT25-T7*gp6.5* | This study |
| YZ1515 | DH5a | pET28b-his.mbp.SUMO | This study |
| YZ1550 | DH5a | pUT18C-*relA*(HS) | This study |
| YZ1551 | DH5a | pUT18C-*relA*(HST) | This study |
| YZ1552 | DH5a | pUT18C-*relA*(TCA) | This study |
| YZ1553 | DH5a | pUT18C-*relA*(CA) | This study |
| YZ1615 | BL21 DE3 | pRARE, pET28b-HMS, pBAD30-T7*gp8*-Flag | This study |
| YZ1620 | BL21 DE3 | pRARE, pET28b-HMS-*relA*, pBAD30-T7*gp8*-Flag | This study |
| YZ1625 | BL21 DE3 | pRARE, pET28b-HMS-*spoT*, pBAD30-T7*gp8*-Flag | This study |
| YZ1632 | BL21 DE3 | pET28a-T7*gp8*-his7 | This study |
| YZ1705 | DH5a | pKT25-*gp8*(Δclip) | This study |
| YZ1706 | DH5a | pKT25-*gp8*(Δ(clip-stem)) | This study |
| YZ1707 | DH5a | pKT25-*gp8*(ΔNt) | This study |
| YZ1708 | DH5a | pKT25-*gp8*(ΔCt) | This study |
| KEIO collection | BW25113 | *trxA::*Kan^R^ | Keio collection^7^ |
| YZ1763 | MG1655 | *trxA::*Kan^R^ | This study |
| YZ1836 | BL21 DE3 | pRARE, pET28b-HMS, pBAD30-T7*gp8*(D220R/E221R)-Flag | This study |
| YZ1837 | BL21 DE3 | pRARE, pET28b-HMS, pBAD30-T7*gp8*(E10K/D11K)-Flag | This study |
| YZ1838 | BL21 DE3 | pRARE, pET28b-HMS-*relA*, pBAD30-T7*gp8*(D220R/E221R)-Flag | This study |
| YZ1839 | BL21 DE3 | pRARE, pET28b-HMS-*relA*, pBAD30-T7*gp8*(E10K/D11K)-Flag | This study |
| YZ1840 | BL21 DE3 | pRARE, pET28b-HMS-*spoT*, pBAD30-T7*gp8*(D220R/E221R)-Flag | This study |
| YZ1841 | BL21 DE3 | pRARE, pET28b-HMS-*spoT*, pBAD30-T7*gp8*(E10K/D11K)-Flag | This study |
| YZ1296 | MG1655 | pBAD30-*gp8*-Flag | This study |
| YZ1857 | MG1655 | pBAD30-*gp8* (D220R/E221R)-Flag | This study |
| YZ1878 | DH5a | pBAD33-*gp59* (N4 portal protein) | This study |
| YZ 1879 | iVEC | pBAD33-P1 portal protein | This study |
| YZ1993 | DH5a | pKT25-*gp8*(D46K/D48K/D53K) | This study |
| YZ1994 | DH5a | pKT25-*gp8*(D99K/D101K) | This study |
| YZ1995 | DH5a | pKT25-*gp8*(D220R/E221R/D222R/E225R) | This study |
| YZ1996 | DH5a | pKT25-*gp8*(D220R/E221R) | This study |
| YZ1997 | DH5a | pKT25-*gp8*(D222R/E225R) | This study |
| YZ1998 | DH5a | pKT25-*gp8*(D241K) | This study |
| YZ1999 | DH5a | pKT25-*gp8*(D457K/D460K) | This study |
| YZ2000 | DH5a | pKT25-*gp8*(D502R) | This study |
| YZ2001 | DH5a | pKT25-*gp8*(E10K/D11K) | This study |
| YZ2002 | DH5a | pKT25-*gp8*(E149K/E151K) | This study |
| YZ2003 | DH5a | pKT25-*gp8*(E192R/D193K) | This study |
| YZ2004 | DH5a | pKT25-*gp8*(E199R/E204R) | This study |
| YZ2005 | DH5a | pKT25-*gp8*(E236R) | This study |
| YZ2006 | DH5a | pKT25-*gp8*(E247R) | This study |
| YZ2007 | DH5a | pKT25-*gp8*(E280R/E284R) | This study |
| YZ2008 | DH5a | pKT25-*gp8*(E416K/E423K) | This study |
| YZ2009 | DH5a | pKT25-*gp8*(E485K/E486K) | This study |
| YZ2010 | DH5a | pKT25-*gp8*(E521R/D528R) | This study |
| YZ2011 | DH5a | pKT25-*gp8*(K4D/R5D) | This study |
| YZ2012 | DH5a | pKT25-*gp8*(K94D) | This study |
| YZ2013 | DH5a | pKT25-*gp8*(K105E) | This study |
| YZ2014 | DH5a | pKT25-*gp8*(K196E) | This study |
| YZ2015 | DH5a | pKT25-*gp8*(K288D/K295D) | This study |
| YZ2016 | DH5a | pKT25-*gp8*(K419D) | This study |
| YZ2017 | DH5a | pKT25-*gp8*(R277D) | This study |
| YZ1794 | MG1655 ∆*trxA* | pCA24N-*gp8*(D220R/E221R) | This study |
| YZ1796 | MG1655 ∆*trxA* | pCA24N-*gp8*(E10KD11K) | This study |
| BP-E-A1 | MG1655 | pBAD33-*gp0.3* | This study |
| BP-E-A3 | MG1655 | pBAD33-*gp1.2* | This study |
| BP-E-A4 | MG1655 | pBAD33-*gp1.6* | This study |
| BP-E-A5 | MG1655 | pBAD33-*gp2.5* | This study |
| BP-E-A6 | MG1655 | pBAD33-*gp3.8* | This study |
| BP-E-A7 | MG1655 | pBAD33-*gp4.2* | This study |
| BP-E-B1 | MG1655 | pBAD33-*gp0.4* | This study |
| BP-E-B2 | MG1655 | pBAD33-*gp0.7* | This study |
| BP-E-B3 | MG1655 | pBAD33-*gp1.3* | This study |
| BP-E-B4 | MG1655 | pBAD33-*gp1.7* | This study |
| BP-E-B5 | MG1655 | pBAD33-*gp2.8* | This study |
| BP-E-B7 | MG1655 | pBAD33-*gp4.3* | This study |
| BP-E-C1 | MG1655 | pBAD33-*gp0.5* | This study |
| BP-E-C3 | MG1655 | pBAD33-*gp1.4* | This study |
| BP-E-C4 | MG1655 | pBAD33-*gp1.8* | This study |
| BP-E-C5 | MG1655 | pBAD33-*gp3* | This study |
| BP-E-C7 | MG1655 | pBAD33-*gp4.5* | This study |
| BP-E-C8 | MG1655 | pBAD33-*gp5.5* | This study |
| BP-E-D1 | MG1655 | pBAD33-*gp0.6A* | This study |
| BP-E-D2 | MG1655 | pBAD33-*gp1.1* | This study |
| BP-E-D3 | MG1655 | pBAD33-*gp1.5* | This study |
| BP-E-D4 | MG1655 | pBAD33-*gp2* | This study |
| BP-E-D5 | MG1655 | pBAD33-*gp3.5* | This study |
| BP-E-D6 | MG1655 | pBAD33-*gp4.1* | This study |
| BP-E-D7 | MG1655 | pBAD33-*gp4.7* | This study |
| BP-E-D8 | MG1655 | pBAD33-*gp5.7* | This study |
| BP-E-E1 | MG1655 | pBAD33-*gp5.9* | This study |
| BP-E-E2 | MG1655 | pBAD33-*gp6.7* | This study |
| BP-E-E3 | MG1655 | pBAD33-*gp8* | This study |
| BP-E-E4 | MG1655 | pBAD33-*gp11* | This study |
| BP-E-E5 | MG1655 | pBAD33-*gp15* | This study |
| BP-E-E6 | MG1655 | pBAD33-*gp18* | This study |
| BP-E-E7 | MG1655 | pBAD33-*gp19.2* | This study |
| BP-E-F1 | MG1655 | pBAD33-*gp6* | This study |
| BP-E-F2 | MG1655 | pBAD33-*gp7* | This study |
| BP-E-F3 | MG1655 | pBAD33-*gp9* | This study |
| BP-E-F4 | MG1655 | pBAD33-*gp12* | This study |
| BP-E-F5 | MG1655 | pBAD33-*gp16* | This study |
| BP-E-F6 | MG1655 | pBAD33-*gp18.5* | This study |
| BP-E-F7 | MG1655 | pBAD33-*gp19.3* | This study |
| BP-E-G1 | MG1655 | pBAD33-*gp6.3* | This study |
| BP-E-G2 | MG1655 | pBAD33-*gp7.3* | This study |
| BP-E-G3 | MG1655 | pBAD33-*gp10A* | This study |
| BP-E-G4 | MG1655 | pBAD33-*gp13* | This study |
| BP-E-G5 | MG1655 | pBAD33-*gp17* | This study |
| BP-E-G6 | MG1655 | pBAD33-*gp18.7* | This study |
| BP-E-G7 | MG1655 | pBAD33-*gp19.5* | This study |
| BP-E-H1 | MG1655 | pBAD33-*gp6.5* | This study |
| BP-E-H2 | MG1655 | pBAD33-*gp7.7* | This study |
| BP-E-H3 | MG1655 | pBAD33-*gp10B* | This study |
| BP-E-H4 | MG1655 | pBAD33-*gp14* | This study |
| BP-E-H5 | MG1655 | pBAD33-*gp17.5* | This study |
| BP-E-H6 | MG1655 | pBAD33-*gp19* | This study |

1	Zhang, Y., Zborníková, E., Rejman, D. & Gerdes, K. Novel (p)ppGpp Binding and Metabolizing Proteins of Escherichia coli. *mBio* **9** (2018). <https://doi.org/10.1128/mBio.02188-17>

2	Kitagawa, M. *et al.* Complete set of ORF clones of Escherichia coli ASKA library (A Complete Set of E. coli K-12 ORF Archive): Unique Resources for Biological Research. *DNA Research* **12**, 291-299 (2006). <https://doi.org/10.1093/dnares/dsi012>

3	Grucela, P. K. & Zhang, Y. E. Basal level of ppGpp coordinates Escherichia coli cell heterogeneity and ampicillin resistance and persistence. *Microb Cell* **10**, 248-260 (2023). <https://doi.org/10.15698/mic2023.11.808>

4	Liu, Y. *et al.* Basal ppGpp regulation by SpoT coordinates metabolic homeostasis and acid resistance. *bioRxiv*, 2026.2001.2026.700336 (2026). <https://doi.org/10.64898/2026.01.26.700336>

5	Gray Michael, J. Inorganic Polyphosphate Accumulation in Escherichia coli Is Regulated by DksA but Not by (p)ppGpp. *Journal of Bacteriology* **201**, 10.1128/jb.00664-00618 (2019). <https://doi.org/10.1128/jb.00664-18>

6	Gray Michael, J. Interactions between DksA and Stress-Responsive Alternative Sigma Factors Control Inorganic Polyphosphate Accumulation in Escherichia coli. *Journal of Bacteriology* **202**, 10.1128/jb.00133-00120 (2020). <https://doi.org/10.1128/jb.00133-20>

7	Baba, T. *et al.* Construction of Escherichia coli K-12 in-frame, single-gene knockout mutants: the Keio collection. *Mol Syst Biol* **2**, 2006.0008 (2006). <https://doi.org/10.1038/msb4100050>
