## Supplemented document 2 for "Phage portal proteins counteract stringent-response–mediated restriction"

**Supplemented document 2:** Plaque formation of wt MG1655 or $\Delta$*relA*$\Delta$*spoT* by 16 different coliphages out of the BASEL collection

| Official phage names and their AlphaFold 3 predicted structures of portal ring | Plaques of infection of MG1655 wt | Plaques of infection of $\Delta$*relA*$\Delta$*spoT* | *$\Delta$relA$\Delta$spoT is more sensitive than wt?* |
|---|---|---|---|
| T5 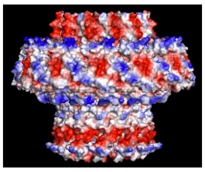 | 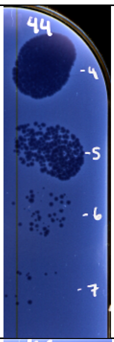  | 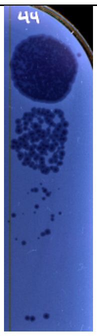  | +/- |
| Lambdavir/HK 97 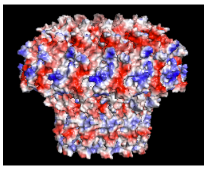 | 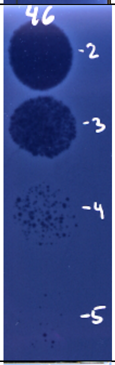  | 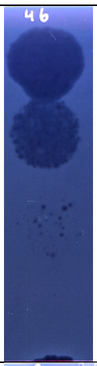  | NO |
| T2 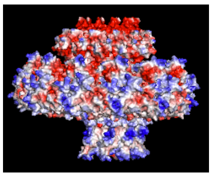 | 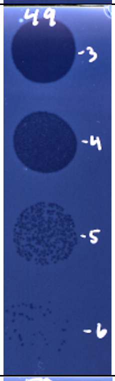  | 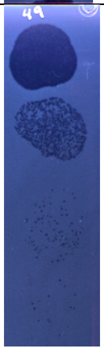  | NO |
| Bas02 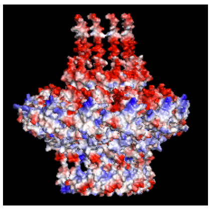 | 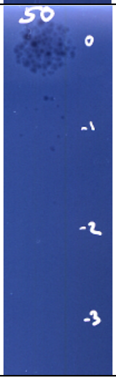  | 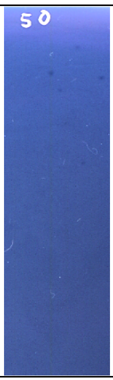  | NO |

| | | | |
|---|---|---|---|
| Bas11 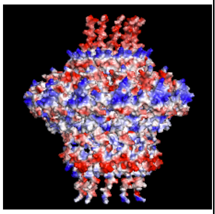 | 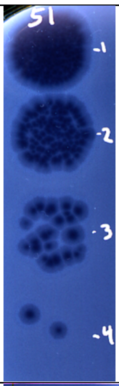  | 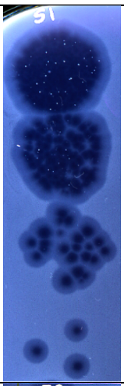  | +/- |
| Bas12 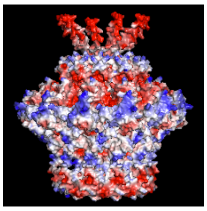 | 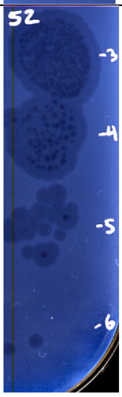  | 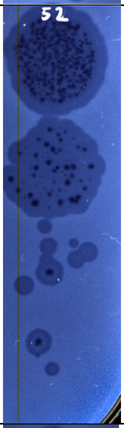  | Yes, darker/clearer plaques |
| Bas17 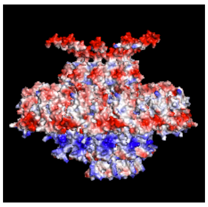 | 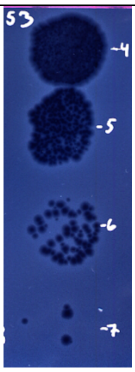  | 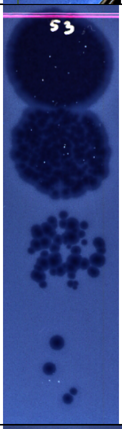  | Yes, bigger and clearer plaques |
| Bas23 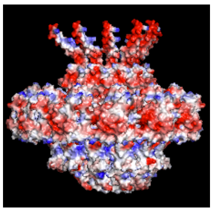 | 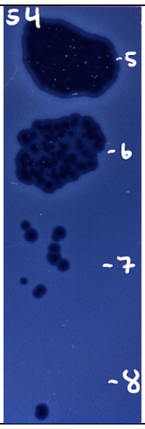  | 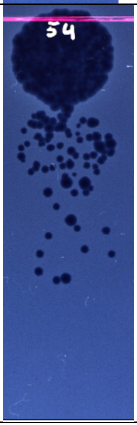  | Yes? darker plaques without halo |

| | | | |
|---|---|---|---|
| Bas26 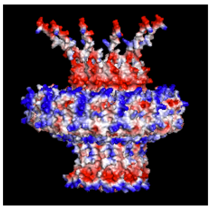 | 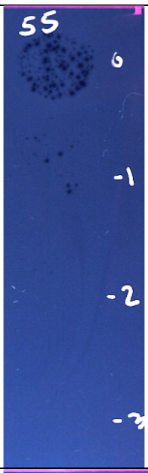 | 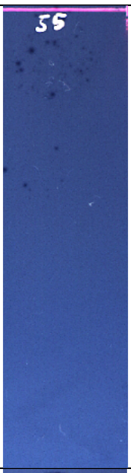 | NO |
| Bas51 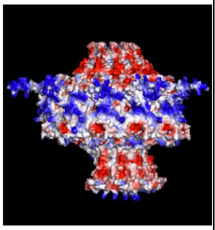 | 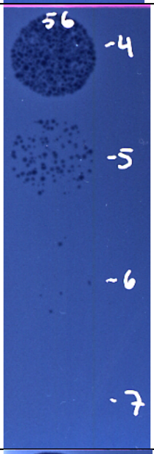 | 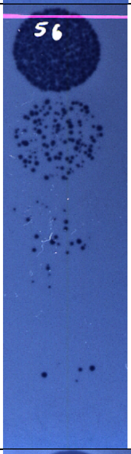 | Yes, more PFU, bigger, darker plaques |
| Bas63 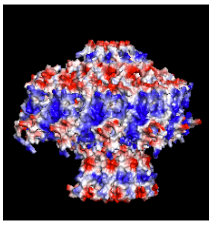 | 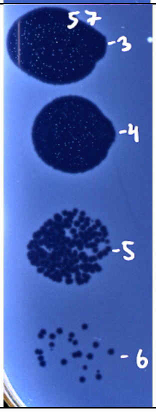 | 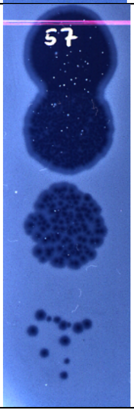 | +/-<br>plaques are bigger but unclear |

| | | | |
|---|---|---|---|
| Bas64 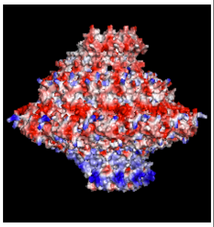 | 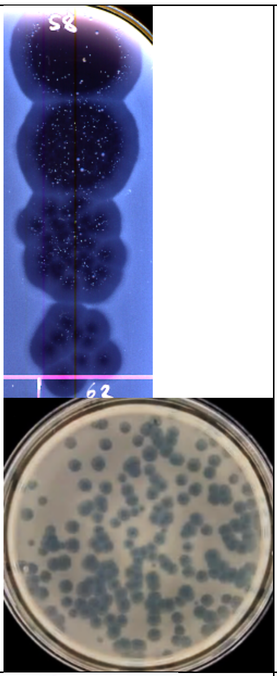 | 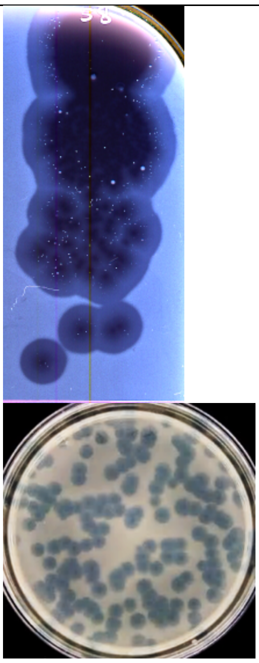 | Yes, bigger plaques, more complete lysis |
| Bas65 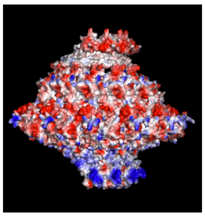 | 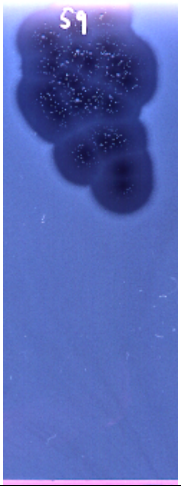 | 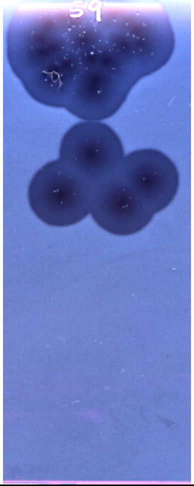 | Yes, bigger plaques |
| Bas66/T3 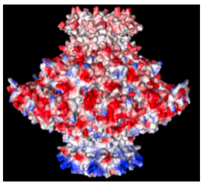 | 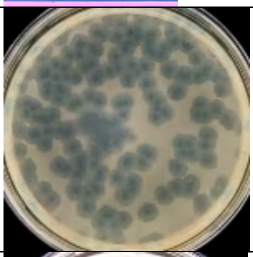 | 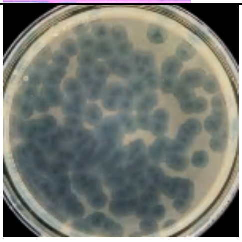 | Yes, bigger and darker plaques |
| Bas68 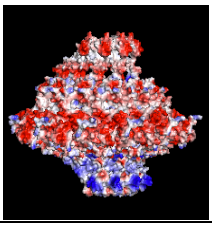 | 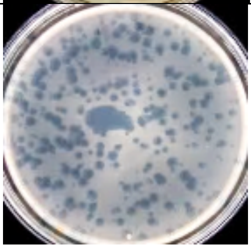 | 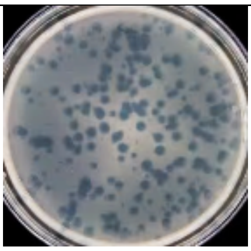 | Yes, darker plaques |

| | | | |
|---|---|---|---|
| Bas69 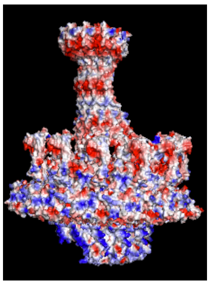 | 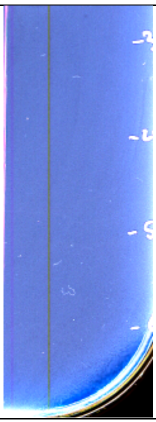 | 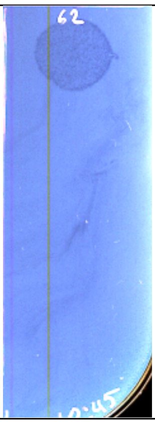 | Yes, more PFU |
