## Supplemented document 3 for "Phage portal proteins counteract stringent-response–mediated restriction"

**Supplemented document 3: all primers used in this study**

| **Primer name or number** | **Description** | **Sequence** |
| --- | --- | --- |
| 0.3-F | Amplify gp0.3 fromT7 and NcoI restriction sequence | CATGccatggCTATGTCTAACATGACTTAC |
| 0.3-R | Amplify gp0.3 fromT7 and EcoRI restriction sequence | gGAATTCttactcttcatcctcctcgta |
| 0.4-F | Amplify gp0.4 fromT7 and with 17 bp flanking sequence for iVEC cloning in pBAD33 | GAAGGAGATATACCATGtctactaccaacgtgc |
| 0.4-R | Amplify gp0.4 fromT7 and 17 bp flanking sequence for iVEC cloning in pBAD33 | cgactctagaggatTTActcagcagattctaaagctattg |
| 0.5-F | Amplify gp0.5fromT7 and NcoI restriction sequence | CATGccatggGTTATATGCTTACTATCGGTCTACTC |
| 0.5-R | Amplify gp0.5 fromT7 and EcoRI restriction sequence | gGAATTCtcatttgcgtagtgcc |
| 0.6A-F | Amplify gp0.6A andgp0.6B fromT7 and NcoI restriction sequence | CATGccatggGTATGAAGCACTACGTTATGC |
| 0.6A-R | Amplify gp0.6A fromT7 and EcoRI restriction sequence | gGAATTCtcagtgcgttctgtcatag |
| 0.7-F | Amplify gp0.7 fromT7 and with 17 bp flanking sequence for iVEC cloning in pBAD33 | GAAGGAGATATACCATGaacattaccgacatcat |
| 0.7-R | Amplify gp0.7 fromT7 and 17 bp flanking sequence for iVEC cloning in pBAD33 | cgactctagaggatTTAgcccattaacattgcg |
| 1.1-F | Amplify gp1.1 fromT7 and NcoI restriction sequence | CATGccatggGTCGTAACTTCGAAAAGATGAC |
| 1.1-R | Amplify gp1.1 fromT7 and EcoRI restriction sequence | gGAATTCttactgaccctcccagctac |
| 1.2-F | Amplify gp1.2 fromT7 and NcoI restriction sequence | CATGccatggGACGTTTATATAGTGGTA |
| 1.2-R | Amplify gp1.2 fromT7 and EcoRI restriction sequence | gGAATTCttacttccagtccttcaact |
| 1.3-F | Amplify gp1.3 fromT7 and NcoI restriction sequence | CATGccatggGTATGAACATTAAGACTAACCCG |
| 1.3-R | Amplify gp1.3fromT7 and EcoRI restriction sequence | gGAATTCttacattttctcttgagggttg |
| 1.4-F | Amplify gp1.4 fromT7 and NcoI restriction sequence | CATGccatggGTTTTAAGAAGGTTGGTAAATTCC |
| 1.4-R | Amplify gp1.4 fromT7 and EcoRI restriction sequence | gGAATTCttaccagttaactatactccacacg |
| 1.5-F | Amplify gp1.5 fromT7 and NcoI restriction sequence | CATGccatggGTATGTACTTAATGCCATTACTCATC |
| 1.5-R | Amplify gp1.5 fromT7 and EcoRI restriction sequence | gGAATTCttaagcgtgaccatctgg |
| 1.6-F | Amplify gp1.6 fromT7 and NcoI restriction sequence | CATGccatggGTTTTCGACTTCATTACAACAAAAG |
| 1.6-R | Amplify gp1.6 fromT7 and EcoRI restriction sequence | gGAATTCtcagaacacctccttgattc |
| 1.7-F | Amplify gp1.7 fromT7 and NcoI restriction sequence | CATGccatggGACTGTTAGATGGTG |
| 1.7-R | Amplify gp1.7fromT7 and EcoRI restriction sequence | gGAATTCttatgcataacatttatccttatg |
| 1.8-F | Amplify gp1.8 fromT7 and NcoI restriction sequence | CATGccatggGTCATAACTTCAAGTCAACCCC |
| 1.8-R | Amplify gp1.8 fromT7 and EcoRI restriction sequence | gGAATTCtcattgacctctcgatttc |
| 2-F | Amplify gp2 fromT7 and NcoI restriction sequence | CATGccatggGTTCAAACGTAAATACAGGTTCAC |
| 2-R | Amplify gp2fromT7 and EcoRI restriction sequence | gGAATTCttacttcggtgctacacaag |
| 2.5-F | Amplify gp2.5 fromT7 and NcoI restriction sequence | CATGccatggCTAAGAAGATTTTCACC |
| 2.5-R | Amplify gp2.5 fromT7 and EcoRI restriction sequence | gGAATTCttagaagtctccgtcttcgtc |
| 2.8-F | Amplify gp2.8 fromT7 and NcoI restriction sequence | CATGccatggAACTGCGGGAGAA |
| 2.8-R | Amplify gp2.8 fromT7 and EcoRI restriction sequence | gGAATTCttagcgccgtaacctg |
| 3-F | Amplify gp3 fromT7 and NcoI restriction sequence | CATGccatggCAGGTTACGGCG |
| 3-R | Amplify gp3 fromT7 and EcoRI restriction sequence | gGAATTCttatttctttcctcctttcctt |
| 3.5-F | Amplify gp3.5 fromT7 and NcoI restriction sequence | CATGccatggCTCGTGTACAGTTTAAAC |
| 3.5-R | Amplify gp3.5 fromT7 and EcoRI restriction sequence | gGAATTCttatccacggtcagaagtg |
| 3.8-F | Amplify gp3.8 fromT7 and NcoI restriction sequence | CATGccatggGTCGCAAGTCTTATAAACAATTCTAT |
| 3.8-R | Amplify gp3.8 fromT7 and EcoRI restriction sequence | gGAATTCctatcggaatcgtgcg |
| 4.1-F | Amplify gp4.1 fromT7 and NcoI restriction sequence | CATGccatggGAACTCGCTGTTCT |
| 4.1-R | Amplify gp4.1 fromT7 and EcoRI restriction sequence | gGAATTCtcattggtttacctcctgag |
| 4.2-F | Amplify gp4.2 fromT7 and NcoI restriction sequence | CATGccatggGTAGGAAGGTCGCCCCGTTT |
| 4.2-R | Amplify gp4.2 fromT7 and EcoRI restriction sequence | gGAATTCttagaatgggactctccagc |
| 4.3-F | Amplify gp4.3 fromT7 and NcoI restriction sequence | CATGccatggGTTTCAAACTGATTAAGAAGTTAGGC |
| 4.3-R | Amplify gp4.3 fromT7 and EcoRI restriction sequence | gGAATTCttactcaaagaatttggaaagc |
| 4.5-F | Amplify gp4.5 fromT7 and NcoI restriction sequence | CATGccatggGTTCTAACGTAGCTGAAACTATCC |
| 4.5-R | Amplify gp4.5 fromT7 and EcoRI restriction sequence | gGAATTCttagttatcaatcttgtcgagg |
| 4.7-F | Amplify gp4.7 fromT7 and NcoI restriction sequence | CATGccatggGTCGTGACCCTAAAGTTATCC |
| 4.7-R | Amplify gp4.7 fromT7 and EcoRI restriction sequence | gGAATTCttatcgtgacttaacaatctcttc |
| 5.5-F | Amplify gp5.5 fromT7 and NcoI restriction sequence | CATGccatggCTATGACAAAGAAATTTAAAG |
| 5.5-R | Amplify gp5.5 fromT7 and EcoRI restriction sequence | gGAATTCtcagaacacctcccgtact |
| 5.7-F | Amplify gp5.7 fromT7 and NcoI restriction sequence | CATGCCATGGGTTCTGACTACCTGAAAGTGCTG |
| 5.7-R | Amplify gp5.7 fromT7 and EcoRI restriction sequence | gGAATTCttagacacatcctcccattc |
| 5.9-F | Amplify gp5.9fromT7 and NcoI restriction sequence | CATGccatggGTTCTCGTGACCTTGTGAC |
| 5.9-R | Amplify gp5.9 fromT7 and EcoRI restriction sequence | gGAATTCtcaagaagtgccattaagtttc |
| 6-F | Amplify gp6.0 fromT7 and NcoI restriction sequence | CATGccatggCACTTCTTGACCT |
| 6-R | Amplify gp6.0 fromT7 and EcoRI restriction sequence | gGAATTCctacggtctccacaggtaaat |
| 6.3-F | Amplify gp6.3 fromT7 and NcoI restriction sequence | CATGccatggAGACCGTAGCGT |
| 6.3-R | Amplify gp6.3 fromT7 and EcoRI restriction sequence | gGAATTCtcacttagtgtcgtgtaacg |
| 6.5-F | Amplify gp6.3 fromT7 and NcoI restriction sequence | CATGccatggGTTTAACACCTATTAACCAATTACTTAAG |
| 6.5-R | Amplify gp6.3 fromT7 and EcoRI restriction sequence | gGAATTCtcaatcctccccatcatc |
| 6.7-F | Amplify gp6.7 fromT7 and NcoI restriction sequence | CATGccatggGTTGTTTCTCACCGAAAATTAAAAC |
| 6.7-R | Amplify gp6.7 fromT7 and EcoRI restriction sequence | gGAATTCtcacttcttacctccaaatg |
| 7-F | Amplify gp7 fromT7 and NcoI restriction sequence | CATGccatggGTTCTGAGTTCACATGTGTGG |
| 7-R | Amplify gp7 fromT7 and EcoRI restriction sequence | gGAATTCttatacctccttaaagtatacacgc |
| 7.3-F | Amplify gp7.3 fromT7 and NcoI restriction sequence | CATGccatggGTAAGAAAGTTAAGAAGG |
| 7.3-R | Amplify gp7.3 fromT7 and EcoRI restriction sequence | gGAATTCttaaatgttgataccgccac |
| 7.7-F | Amplify gp7.7 fromT7 and NcoI restriction sequence | CATGccatggAAGACTGCATTGAATG |
| 7.7-R | Amplify gp7.7 fromT7 and EcoRI restriction sequence | gGAATTCttaaatgtgtctccatgtcttac |
| 8-F | Amplify gp8 fromT7 and NcoI restriction sequence | CATGccatggCTGAGAAACGAAC |
| 8-R | Amplify gp8 fromT7 and EcoRI restriction sequence | gGAATTCttaaattcccggctgtaaac |
| 9-F | Amplify gp9 fromT7 and NcoI restriction sequence | CATGccatggCTGAATCTAATGCAGAC |
| 9-R | Amplify gp9 fromT7 and EcoRI restriction sequence | gGAATTCtcagaagttcgaatcgattac |
| 10A-F | Amplify gp10A and gp10B fromT7 and NcoI restriction sequence | CATGccatggCTAGCATGACTGGT |
| 10A-R | Amplify gp10A fromT7 and EcoRI restriction sequence | gGAATTCttactccactttgaaaaccac |
| 10B-R | Amplify gp10B fromT7 and EcoRI restriction sequence | gGAATTCttattgctcagcggtg |
| 11-F | Amplify gp11 fromT7 and NcoI restriction sequence | CATGccatggGTCGCTCATACGATATGAACG |
| 11-R | Amplify gp11 fromT7 and EcoRI restriction sequence | gGAATTCttagcgagtcagtagaccag |
| 12-F | Amplify gp12 fromT7 and NcoI restriction sequence | CATGccatggCACTCATTAGCC |
| 12-R | Amplify gp12 fromT7 and EcoRI restriction sequence | gGAATTCttaaataccggaacttctcc |
| 13-F | Amplify gp13 fromT7 and NcoI restriction sequence | CATGccatggGTATGACTATAAGACCTACTAAAAGTACAG |
| 13-R | Amplify gp13 fromT7 and EcoRI restriction sequence | gGAATTCttatcctcctttcgtgattg |
| 14-F | Amplify gp14 fromT7 and NcoI restriction sequence | CATGccatggGTTGTTGGGCAGCCGCAATAC |
| 14-R | Amplify gp14 fromT7 and EcoRI restriction sequence | gGAATTCttacctccccgtcttg |
| 15-F | Amplify gp15 fromT7 and with 21bp flanking sequence for iVEC cloning in pET28b | TAAGAAGGAGATATACCATGGgtagtaaaattgaatctgcccttc |
| 15-R | Amplify gp15 fromT7 and 21 bp flanking sequence or iVEC cloning in pET28b | ttgtcgacggagctcgaattcTTACTCCTTACGTCCGTAGATG |
| 16-F | Amplify gp16 fromT7 and NcoI restriction sequence | CATGccatggATAAGTACGATAAGAACG |
| 16-R | Amplify gp16 fromT7 and EcoRI restriction sequence | gGAATTCttatttcctacgctccctc |
| 17-F | Amplify gp17 fromT7 and NcoI restriction sequence | CATGccatggCTAACGTAATTAAAACC |
| 17-R | Amplify gp17 fromT7 and EcoRI restriction sequence | gGAATTCttactcgttctccaccat |
| 17.5-F | Amplify gp17.5 fromT7 and NcoI restriction sequence | CATGccatggGTCTATCATTAGACTTTAACAACGAAT |
| 17.5-R | Amplify gp17.5 fromT7 and EcoRI restriction sequence | gGAATTCtcactccttattggctttct |
| 18-F | Amplify gp18 fromT7 and NcoI restriction sequence | CATGccatggAAAAGGATAAGAGCC |
| 18-R | Amplify gp18 fromT7 and EcoRI restriction sequence | gGAATTCtcactgtaatgtgtaaatatcatc |
| 18.5-F | Amplify gp18.5 fromT7 and NcoI restriction sequence | CATGccatggGTCTAGAATTTTTACGTAAGCTAATCC |
| 18.5-R | Amplify gp18.5 fromT7 and EcoRI restriction sequence | gGAATTCctacttacgttgcagttcac |
| 18.7-F | Amplify gp18.7 fromT7 and NcoI restriction sequence | CATGccatggGTAGTACGTTAAGAGAGTTGAGGCT |
| 18.7-R | Amplify gp18.7 fromT7 and EcoRI restriction sequence | gGAATTCtcactgcgagaatacgtttag |
| 19-F | Amplify gp19 fromT7 and NcoI restriction sequence | CATGccatggGTTCTACTCAATCCAATCGTAATG |
| 19-R | Amplify gp19 fromT7 and EcoRI restriction sequence | gGAATTCtcaccactcaatgaaagac |
| 19.2-F | Amplify gp19.2 fromT7 and NcoI restriction sequence | CATGccatggGTACACAACCATTATCTG |
| 19.2-R | Amplify gp19.2 fromT7 and EcoRI restriction sequence | gGAATTCtcagcgggtacttctcg |
| 19.3-F | Amplify gp19.3 fromT7 and NcoI restriction sequence | CATGccatggCTACTCCGATAAGACC |
| 19.3-R | Amplify gp19.3 fromT7 and EcoRI restriction sequence | gGAATTCtcataccacgggcac |
| 19.5-F | Amplify gp19.5 fromT7 and NcoI restriction sequence | CATGccatggGTTTCCGCTTATTGTTGAAC |
| 19.5-R | Amplify gp19.5 fromT7 and EcoRI restriction sequence | gGAATTCctaatcgctacaagtgagtatagag |
| pYZ473 | XbaI-spoTNt-F | gCTCTAGAGCcttatctgtttg |
| pYZ474 | XmaI-spoTSyn-R | tcccCCCGGGgTTACGGGAAGAGATCGGA |
| pYZ475 | XmaI-spoTtgs-R | tcccCCCGGGgTTACGGAGCGGTAATGATTTC |
| pYZ476 | XmaI-spoTCt-R | tcccCCCGGGGTTAATTTCG |
| pYZ477 | XbaI-spoTtgs-F | gcTCTAGAGCctGATCTCTTCCCGGATGAG |
| pYZ478 | XbaI-spoTcc-F | gcTCTAGAGCctGGCGCTCGCCCGAAT |
| pYZ479 | XbaI-Spot-F | gcTCTAGAGCCTTATCTGTTTGAAAGCCTG |
| pYZ480 | XmaI-Spot-R | tcccCCCGGGGTTAATTTCGGTTTCGGGTG |
| pYZ481 | XbaI-relA-F | gcTCTAGAGCCTGTTGCGGTAAGAAGTG |
| pYZ482 | XmaI-relA-R | tcccCCCGGGATTTAACTCCCGTGCAACC |
| pYZ684 | EcoRI-SUMO-F | gGAATTCggttcggactcagaagtcaa |
| pYZ685 | EcoRI-SUMO-R | gGAATTCtaataccgtcgtacaagaat |
| pYZ692 | NcoI-His-sumo-F | ggccatggttcggactcagaagtcaatc |
| pYZ693 | sumo-spoT-R | ttggtggttatctgtttgaaagcctgaatcaac |
| pYZ694 | sumo-spoT-F | aacagataaccaccaatctgttctctgtgag |
| pYZ695 | HindIII-spoT-R | ccgcaagcTTAATTTCGGTTTCGGGTGACTTTAATC |
| pYZ702 | spoT-iVEC-cloning-F1 | CAGGCGTATCTCGTTGCAC |
| pYZ703 | spoT-iVEC-cloning-R1 | CACTGCATAAGCGAAGTCG |
| pYZ862 | QC-rmvNcoI-pkt25-R | CGTGACCgATGGCGATGCCTGCT |
| pYZ863 | QC-rmvNcoI-pkt25-F | TCGCCATcGGTCACGACGAGATCCTC |
| pYZ864 | QC-insNcoI-B2H-R | GAATTCccatgGGTACCCGGGGATCC |
| pYZ865 | QC-insNcoI-pKT-F | GTACCcatggGAATTCACTGGCCGTCG |
| pYZ866 | QC-insNcoI-pUT-F | GTACCcatggGAATTCATCGATATAACTAAGTAATATGGTG |
| pYZ867 | HindIII-rpoZ-F | CCCaagcttaGCACGCGTAACTGTTCAG |
| pYZ868 | XbaI-rpoZ-R | GCtctagactACGACGACCTTCAGCAATA |
| pYZ869 | pK(N)T25-seq-R | ccttcgccacggccttgatg |
| pYZ870 | pK(N)T25-seq-F | TTCCGGCTCGTATGTTG |
| pYZ871 | QC-respKNT25-R | GAATTCGAGCTCGGTACCCGGGGATCCTCTAGAGTCGACCTGCAGGCATGCAAGCTTGGCGTAATCATGGTCATAGC |
| pYZ872 | QC-respKNT25-F | GTACCGAGCTCGAATTCAATGACCATGCAGCAATCG |
| pYZ905 | QCpET-HMS-BamHI-F | tggtGGATCCTAAGCTTGCGGCCGC |
| pYZ906 | QCpET-HMS-BamHI-R | agcttAGGATCCACCAATCTGTTCTCTGTG |
| pYZ917 | XbaI-relApUT18C-F | gcTCTAGAGCctGTTGCGGTA |
| pYZ918 | XmaI-relAHST-R | TCCCcccgggaATCTTTACTGCGGTTTTGCG |
| pYZ919 | XmaI-relAHS-R | TCCCcccgggaGTCAAAGACCTGACTACGTACTTC |
| pYZ920 | XmaI-relApUT18C-R | TCCCcccgggaACTCCCGTGCAACCG |
| pYZ921 | XbaI-relA(TCA)-F | gcTCTAGAGCctGTCTTTGACGACCGGGT |
| pYZ922 | XbaI-relA(CA)-F | gcTCTAGAGCctGACAAAAACATTCTGGCTGG |
| pYZ927 | BamHI-relA-F | CGggatccGTTGCGGTAAGAAGTGCA |
| pYZ928 | HindIII-relA-R | CCCaagcttaACTCCCGTGCAACC |
| pYZ1063 | *gp8*(clip deltion)-R | TCCAGTTGGCCGATAACCTTAGAGCTAATCA |
| pYZ1064 | *gp8*(clip deltion)-F | ttatcggcCAACTGGAGAAGCAAGCAG |
| pYZ1065 | *gp8*(clip and stem deltion)-R | AACATAAAgtacgaacgaccgtaggatt |
| pYZ1066 | *gp8*(clip and stem deltion)-F | gttcgtacTTTATGTTGAACTCTGCGGTT |
| pYZ1067 | *gp8*(N-terminus deletion)-R | TTAGGGAACATGGGTACCCGGGG |
| pYZ1068 | *gp8*(N-terminus deletion)-F | TACCcatgTTCCCTAAGGACTCCGATAAC |
| pYZ1069 | *gp8*(C-terminus deletion)-R | aattcTTAggtgagtagaataccagaagtgtc |
| pYZ1070 | *gp8*(C-terminus deletion)-F | tactcaccTAAgaattcACTggccgtc |
| pYZ1187 | Confirm deletion of trxA-F | CGAAAGCGTATCCGGTG |
| pYZ1188 | Confirm deletion of trxA-R | GCGTCCAGTTTTTAGCGAC |
| pYZ1189 | Amplify trxA from MG1655 with 40 bp gp8 flanking sequence-F | taacgcgaatactacagcgtaagacatggagacacatttaatgagcgataaaattattcacc |
| pYZ1190 | Amplify trxA from MG1655 with 40 bp gp8 flanking sequence-R | cgctcatttcaaagatgaggtctccctatagtgagtcgtattacgccaggttagcg |
| pYZ1191 | Amplify gp8+350 bp flanking region and XhoI restriction sequence-F | CCGctcgagGGTTATGGTCGTAAGTGG |
| pYZ1192 | Amplify gp8+350 bp flanking region and HindIII restriction sequence-R | CCCaagcttCTCAGAGCCATCACCG |
| pYZ1193 | Confirm gp8 in cPCR-R | TCTTGTCCTCGACCAATTG |
| pYZ1194 | Confirm gp8 in cPCR-F | TATCTGGATGAGGACTCAGG |
| pYZ1195 | Confirm trxA gene in T7-F | AGCGTAAGACATGGAGACAC |
| pYZ1196 | Confirm trxA gene in T7-R | TCATTTCAAAGATGAGGTCTC |
| pYZ34 | Confirm gp8 sequence on pCA24N-F | GGCCCTTTCGTCTTCACCTC |
| pYZ35 | Confirm gp8 sequence on pCA24N-R | GGCAACCGAGCGTTCTGAAC |
| pYZ1085 | QC of gp8-D46KD48KD53K-F | AAGTCCAAGAACGCCTCTACAAAGTATCAAACTCCGTGGCAAG |
| pYZ1086 | QC of gp8-D46KD48KD53K-R | CTTTGTAGAGGCGTTCTTGGACTTCTTAGGGAACAATGATGGGA |
| pYZ1087 | QC of gp8-D99K/D101K-F | TGAGCAAACCCAAAGGACTCGCTAAGGTCGATG |
| pYZ1088 | QC of gp8-D99K/D101K-R | AGTCCTTTGGGTTTGCTCAGTAACTGCTTTGCTTC |
| pYZ1089 | QC of gp8-D220R/E221R/D222R/E225R-F | AGACGTAGGTCAGGTAGGTACCTCCGATACGAAGAGGT |
| pYZ1090 | QC of gp8-D220R/E221R/D222R/E225R-R | CCTACCTGACCTACGTCTCAGATAGATGTGAGTGTACACGTC |
| pYZ1091 | QC of gp8-D220R/E221R-F | ATCTGCGACGTGACTCAGGTGAATACCTCCG |
| pYZ1092 | QC of gp8-D220R/E221R-R | GAGTCACGTCGCAGATAGATGTGAGTGTACACGTC |
| pYZ1093 | QC of gp8-D222R/E225R-F | AGCGTTCAGGTAGGTACCTCCGATACGAAGAGGT |
| pYZ1094 | QC of gp8-D222R/E225R-R | TACCTACCTGAACGCTCATCCAGATAGATGTGAGTGTA |
| pYZ1095 | QC of gp8-D241K-F | AGGCTCCAAAGGGACTTATCCTAAAGAGGCTT |
| pYZ1096 | QC of gp8-D241K-R | AAGTCCCTTTGGAGCCTTGGACTTCCA |
| pYZ1097 | QC of gp8-D457D460KK-F | GGAAAGACCCTAAGATTAACCTTGCGATGATTAAGTTAC |
| pYZ1098 | QC of gp8-D457D460KK-R | ATCTTAGGGTCTTTCCGCATAGGTGCCAGT |
| pYZ1099 | QC of gp8-D502R-F | GTATGAGGAATGGTGCTGCTGCG |
| pYZ1100 | QC of gp8-D502R-R | CACCATTCCTCATACCCATTTGCATAGACTGT |
| pYZ1101 | QC of gp8-E10KD11K-F | TTGCGAAGAAAGGCGCAAAGTCTGTCTATG |
| pYZ1102 | QC of gp8-E10KD11K-R | GCGCCTTTCTTCGCAAGTCCTGTTCGTTT |
| pYZ1103 | QC of gp8-E149K/E151K-F | ACCGAAGCCGAAGGGGTCAAACTATAATCCCATGA |
| pYZ1104 | QC of gp8-E149K/E151K-R | ACCCCTTCGGCTTCGGTAGGTACAGCAGGAC |
| pYZ1105 | QC of gp8-E192R/D193K-F | CTAGAAAGATCCGTAAGGCTGTAGAAGGT |
| pYZ1106 | QC of gp8-E192R/D193K-R | TACGGATCTTTCTAGGGAGAGCACCAAAAGC |
| pYZ1107 | QC of gp8-E199R/E204R-F | AGGGGTCAAGGTGGTAGAAAGAAAGCTGATGAGACAATCG |
| pYZ1108 | QC of gp8-E199R/E204R-R | TCTACCACCTTGACCCCTTACAGCCTTACGGATGTCCTC |
| pYZ1109 | QC of gp8-E236R-F | GGGTATGAGAGTCCAAGGCTCCGATG |
| pYZ1110 | QC of gp8-E236R-R | TTGGACTCTCATACCCTCGACCTCTTCG |
| pYZ1111 | QC of gp8-E247R-F | TCCTAAAAGGGCTTGCCCATACATCC |
| pYZ1112 | QC of gp8-E247R-R | GCAAGCCCTTTTAGGATAAGTCCCATCGG |
| pYZ1113 | QC of gp8-E280R/E284R-F | TCGCAATCTCCAACGCGCTATCGTCAAGATGTCCAT |
| pYZ1114 | QC of gp8-E280R/E284R-R | CGCGTTGGAGATTGCGAAGGGACCGTAAGTCACCTA |
| pYZ1115 | QC of gp8-E416KE423K-F | AAGTTACCTAAGGAAGCCGTAAAGCCAACCATTAGTACAGGTCTG |
| pYZ1116 | QC of gp8-E416KE423K-R | TTACGGCTTCCTTAGGTAACTTAGGAATCTGTTGCGTGGC |
| pYZ1117 | QC of gp8-E485E486KK-F | TCACCAAGAAGCAGAAGCAACAGAAGATGGC |
| pYZ1118 | QC of gp8-E485E486KK-R | TTCTGCTTCTTGGTGAGTAGAATACCAGAAGTGTCA |
| pYZ1119 | QC of gp8-E521R/D528R-F | CGAGCTATGGCTGCTGCCGCTCGATCCGTAGGTTTACAGCCG |
| pYZ1120 | QC of gp8-E521R/D528R-R | TCGAGCGGCAGCAGCCATAGCTCGAGGTGAAGCTGTAGCTTGTG |
| pYZ1121 | QC of gp8-K4D/R5D-F | CTGAGGACGACACAGGACTTGCGGAGGA |
| pYZ1122 | QC of gp8-K4D/R5D-R | CCTGTGTCGTCCTCAGCCATGGGTACCC |
| pYZ1123 | QC of gp8-K94D-F | TGAAGCAGACCAGTTACTGAGCGACCCC |
| pYZ1124 | QC of gp8-K94D-R | AACTGGTCTGCTTCATATTCAGATATAGTAAGTCG |
| pYZ1125 | QC of gp8-K105E-F | ACTCGCTGAGGTCGATGAGGGCCT |
| pYZ1126 | QC of gp8-K105E-R | TCGACCTCAGCGAGTCCATCGGG |
| pYZ1127 | QC of gp8-K196E-F | CATCCGTGAGGCTGTAGAAGGTCAAGGTG |
| pYZ1128 | QC of gp8-K196E-R | ACAGCCTCACGGATGTCCTCAGGGAG |
| pYZ1131 | QC of gp8-K288D/K295D-F | GATATGTCCATGATTAGCTCTGATGTTATCGGCTTAGTGAATCCTG |
| pYZ1132 | QC of gp8-K288D/K295D-R | ATCAGAGCTAATCATGGACATATCGACGATAGCCTCTTGGAGATT |
| pYZ1135 | QC of gp8-R277D-F | GTGACTTAGACTCCCTTGAAAATCTCCAAG |
| pYZ1136 | QC of gp8-R277D-R | AGGGAGTCTAAGTCACCTAAGTATTCCTCAATGTA |
| pYZ1318 | iVEC-N4-gp59-F2 | gaattcAAGAAGGAGATATACCatgGAACAAAACACTGATTCTATGGTTC |
| pYZ1319 | iVEC-N4-gp59-R2 | TCTCATCCGCCAAAACAGCCAAGCTTAATTACCTAATCTAATTCCTGGGT |
| pYZ1320 | iVEC-P1-portalp-F | gaattcAAGAAGGAGATATACCatgGCAGACAATAAAATCACGCTATC |
| pYZ1321 | iVEC-P1-portalp-R | CTCATCCGCCAAAACAGCCAAGCTTAGTCATTATCGTTTCCCTCTTT |
| pYZ1285 | iVEC-P1-portalp-F | GTCGACTCTAGAGGATCCCCGGGTAGCAGACAATAAAATCACGCTATC |
| pYZ1286 | iVEC-P1-portalp-R | GTTGtaaaacgacggccAGTgaatTCAGTCATTATCGTTTCCCTC |
| pYZ1287 | iVEC-N4-gp59-F | GTCGACTCTAGAGGATCCCCGGGTAGAACAAAACACTGATTCTATGGTTC |
| pYZ1288 | iVEC-N4-gp59-R | GTTGtaaaacgacggccAGTgaaTTAATTACCTAATCTAATTCCTGGGT |
| pYZ1293 | N4gp59-seqF | GTTCCTTTAGCAAACCACC |
| pYZ1294 | P1portalP-seqF | GGCCTAAGTCAAACCTTATGC |
